# Scaling and ecomorphology of lagomorph body shape and appendicular skeleton

**DOI:** 10.64898/2026.05.07.723560

**Authors:** Coby Huizenga, Nia Brice, Chris J. Law

## Abstract

The diversity of body shapes is one of the most prominent features of phenotypic variation in mammals. Yet, mammalian body shapes are poorly quantified and the underlying components contributing to its diversity as well as its relationship to other components of the skeleton are rarely tested. Here, we use lagomorphs (hares, rabbits and pikas) as a model system to (1) investigate which components of the skeleton contributed the most to body shape diversity, (2) examine the relationships between body shape and relative limb lengths, and (3) test how body size, ecotype, burrowing behavior, and locomotor mode influenced variation in lagomorph body shape and appendicular morphology. We quantified the body shape and functional proxies of the appendicular skeleton in 40 lagomorph species from osteological specimens held at museum collections. Using phylogenetic comparative methods, we found the relative length of the ribs and elongation or shortening of the thoracic and lumbar regions contributed the most to body shape evolution across lagomorphs. Second, we found that only leporids (hares and rabbits) exhibited a significant relationship between limb length and body shape, where more elongate species exhibit relatively shorter forelimbs and hindlimbs. Lastly, we found that models incorporating body size were the best predictors of lagomorph body shape and the majority of the appendicular traits, whereas models incorporating burrowing behavior and locomotor mode were largely poor fits. Broadly, these results indicate that larger lagomorphs tend to exhibit more robust body shapes with longer, more gracile forelimbs, whereas smaller lagomorphs tend to exhibit more elongate body shapes with shorter, more robust forelimbs. Overall, this work contributes to the growing understanding of mammalian body shape evolution and demonstrates the importance of not omitting body size in ecomorphological analyses.

## Introduction

Understanding the patterns of morphological diversity found across the tree of life is a major goal of organismal biology. Several factors have been identified as contributors to morphological diversity such as allometry, ecology, behavior, and phylogenetic history, the last of which further provides the framework for understanding its evolution (e.g., Goswami et al. 2022; Guillerme et al. 2023; Grossnickle et al. 2024; Law et al. 2024; Title et al. 2024; Hodge et al. 2025). Size is often hypothesized to be a line of least evolutionary resistance for morphological evolution (Schluter 1996; Marroig and Cheverud 2005, 2010); thus, scaling is often found to be a strong predictor of variation in many morphological traits (e.g., Bertram and Biewener 1990; Martín-Serra et al. 2014; Law 2021b; Holding et al. 2022; Mitchell et al. 2024). However, evolutionary changes in size may be constrained, resulting in species evolving distinct shapes or proportions of traits (i.e., size-independent morphologies) as adaptations towards specific ecological functions (Zelditch et al. 2017; Mitchell et al. 2023). As a result, evolution of morphological traits that deviate from predicted scaling patterns may lead to morphological specializations or diversification in some species or entire clades (Collar et al. 2011; Law et al. 2018; Friedman et al. 2019; Arbour et al. 2021; Zelditch and Swiderski 2022; Mitchell et al. 2023).

Mammals are of particular interest for examining morphological variation due to their apparent extremes in body sizes and body plans, ranging from small elongate, short-legged weasels to long-legged ungulates to large, robust elephants. These distinct body plans often reflect the diversity of their habitats, allowing them to run, swim, swing, and/or dig through their environment. Impressively, mammals exhibit these diverse body plans despite conserved skeletal elements, particularly in the vertebral column (Galis 1999; Narita and Kuratani 2005; Young and Hallgrímsson 2005; Jones et al. 2020; Taewcharoen et al. 2024). The evolution of the mammalian body shape is typically described along a continuum of elongation and robustness (Collar et al. 2013; Law 2021b; Linden et al. 2023). Because vertebral number is constrained in most mammals (Narita and Kuratani 2005), evolutionary changes of mammalian body shape along this continuum occur through relative lengthening or shortening of the head or vertebrae and/or changes in body depth rather than changes in the number of vertebrae observed in many non-mammalian vertebrates (Collar et al. 2013; Law 2021b; Linden et al. 2023). Thus, despite constraints on vertebral number, multiple diverging evolutionary pathways have led to the diversity of mammalian body shape we see today (Law 2021b, 2022). Although evolutionary allometry strongly influences the evolution of body shape diversity, the direction of allometry is dependent on ecological factors (Law 2021a; Linden et al. 2023). For instance, terrestrial carnivorans and ground squirrels exhibit negative body shape allometry, in which larger species exhibit more robust bodies whereas smaller species exhibit more elongate bodies. In contrast, fully aquatic carnivorans (i.e., pinnipeds) and gliding squirrels exhibit the opposite trend in which larger species exhibit more elongate bodies whereas smaller species exhibit more robust bodies (Law 2021a; Linden et al. 2023). Similarly, body size, ecological factors, and evolutionary history contribute to the diversity of the appendicular skeleton (e.g., Kilbourne and Hoffman 2013; Martín-Serra et al. 2014; Wölfer et al. 2019; Etienne et al. 2021; Burtner et al. 2024). Within many mammalian clades, the gradient from gracility to robustness in limb bones is a major trend found in the diversity of the appendicular skeleton, and is generally associated with a functional trade-off between increasing cost of transport associated with cursoriality and resisting stresses associated with locomoting through resistant media (Samuels and Van Valkenburgh 2008; Martín-Serra et al. 2014; Kilbourne 2017; Hedrick et al. 2020; Muñoz 2020; Marshall et al. 2021; Rickman et al. 2023; Law et al. 2025). However, the appendicular skeleton is also influenced by size because more mechanical support is needed to compensate for an increase in the mechanical forces exerted during locomotion at larger body sizes (McMahon 1973; Alexander et al. 1979; Bertram and Biewener 1990; Christiansen 1999). Thus, scaling has strong effects on the relationship between limb morphologies and locomotor mode (Scheidt et al. 2019; Wölfer et al. 2019; Rickman et al. 2023) and may even be the primary force contributing to limb diversity rather than ecological factors (Rothier et al. 2025).

Although the mammalian body plan is clearly diverse, few studies have simultaneously examined whether body shape and the limbs are similarly affected by size, ecological factors, and evolutionary history. In this study, we investigated how these factors influenced various aspects of mammalian body shape (defined by the head-body elongation ratio (Law et al. 2019)) and appendicular skeleton in lagomorphs. Lagomorphs are a great model system to explore diversity in the body plan due to their morphological, behavioral, and size variation. Furthermore, lagomorphs exhibit unique Lagomorphs consist of approximately 109 species that are often categorized into three ecotypes: pikas, rabbits, and hares (Kraatz et al. 2021). Locomotory modes vary among these ecotypes with hares, rabbits, and pikas typically exhibiting cursorial, saltatorial, and scrambling behaviors, respectively (Wilson et al. 2016; Kraatz et al. 2021). Despite their diverse locomotor behaviors and high species richness, macroevolutionary analyses of the lagomorph skeletal system have received little attention except in the cranium (Ge et al. 2015; Kraatz et al. 2015; Kraatz and Sherratt 2016; Wood Bailey et al. 2025). In the appendicular skeleton, researchers have found that increasing cursoriality gradient from pikas to rabbits to hares is associated with more gracile proximal limb bones for increased stride length whereas the distal bones remain relatively robust to resist fracture from impact (Camp and Borell 1937; Gambaryan 1974; Young et al. 2014). However, these studies often only use a single species as a representative of each ecotype. Just doubling the number of species per ecotype (i.e., from one species to two) weakened this limb-cursoriality relationship, possibly due to variation from body size and burrowing behavior (Martin et al. 2022). Although lagomorphs exhibit a broad range of burrowing behaviors from species that dig complex forms to species that do not burrow at all (Wilson et al. 2016), how burrowing behaviors are associated with limb morphology remains to be investigated in this group. In contrast to the skull and limbs, morphological diversity in the rest of the lagomorph body plan such as body shape or axial skeleton has not been investigated across the clade; thus, how size and ecological and behavioral factors influence these aspects of morphological diversity also remains unknown in lagomorphs.

The goals of this study were three-fold. First, we created a dataset that quantifies the body shapes of lagomorphs. We then tested which cranial and axial components contributed the most to overall body shape diversity as well as whether body shape diversity is associated with evolutionary changes in the lengths of the forelimb and hindlimb across lagomorphs. Third, we assessed whether body size, ecotype, locomotory mode, and/or burrowing behavior best predicted body shape diversity and limb ecomorphology across lagomorphs. We predicted that body size will have the strongest influence on body shape diversity as previous work indicated that increased body size is associated with increased body robustness in terrestrial mammals (Law et al. 2019; Law 2021b; Linden et al. 2023). In the limbs, we predicted that both burrowing behavior and cursoriality would have the strongest effect on limb ecomorphology. Increased burrowing behavior would be associated with more robust limbs to accommodate increased muscle attachments and greater mechanical advantage for greater output force during digging whereas cursoriality would be associated with more elongated limbs to support the longer strides taken during cursorial locomotion.

## Methods

### Morphological Data

We acquired 70 adult skeletal specimens of 40 lagomorph species from eight natural history museums. Adults were determined by fusion of sutures on the vertebrae and limb bones. We quantified the body shape of each specimen using the head-body elongation ratio (hbER) (Law et al. 2019), which was calculated as the sum of the length of the head and body divided by body depth ((head length + body length) / body depth). Head length was measured as the condylobasal length of the cranium from the anteriormost point on the premaxilla to the posteriormost point on the surfaces of the occipital condyles; body length was measured by summing the centrum lengths (measured along the ventral surface of the vertebral centrum) of each cervical, thoracic, lumbar, and sacral vertebrae; and body depth was estimated as the average length of the four longest ribs. Eac h rib was measured as a curve from the end of the capitulum to the point of articulation with the costal cartilage using a flexible measuring tape. High values of hbER suggest a more elongate body shape whereas low values of hbER suggest a more robust body shape.

We further examined how the cranial and axial skeleton contributed to overall body shape. We quantified head elongation ratio (headER) by dividing cranial length with cranial height. For each vertebral region (V; i.e., cervical, thoracic, lumbar, and sacral), we calculated the axial elongation index (AEI_V_) as the total sum of vertebral lengths (L_V_) divided by the average vertebral height (L_H_; measured from the ventral surface of the centrum to the tip of the neural spine): AEI_V_ = L_V_ /mean(H_V_). We also quantified body size using the geometric mean of the cranial, vertebral, and rib measurements (Nth root of the product of our measurements for the cranium, vertebrae, and ribs; N = 11) (Mosimann 1970; Klingenberg 2016).

We then measured 11 forelimb (scapula, humerus, radius, ulna and third metacarpal) and 10 hindlimb (femur, tibia, calcaneus and third metatarsal) traits (Fig. 1). From these traits, we calculated the forelimb length as the sum of the lengths of the scapula, humerus, radius, and third metacarpal and the hindlimb length as the sum of the lengths of the femur, tibia, and third metatarsal. We also calculated 13 morphological proxies of function as proxies for functional traits (Table 1). These functional proxies are often used to capture the functional diversity of the limbs (Davis 1964; Samuels and Van Valkenburgh 2008; Law et al. 2025).

**Fig. 1.**
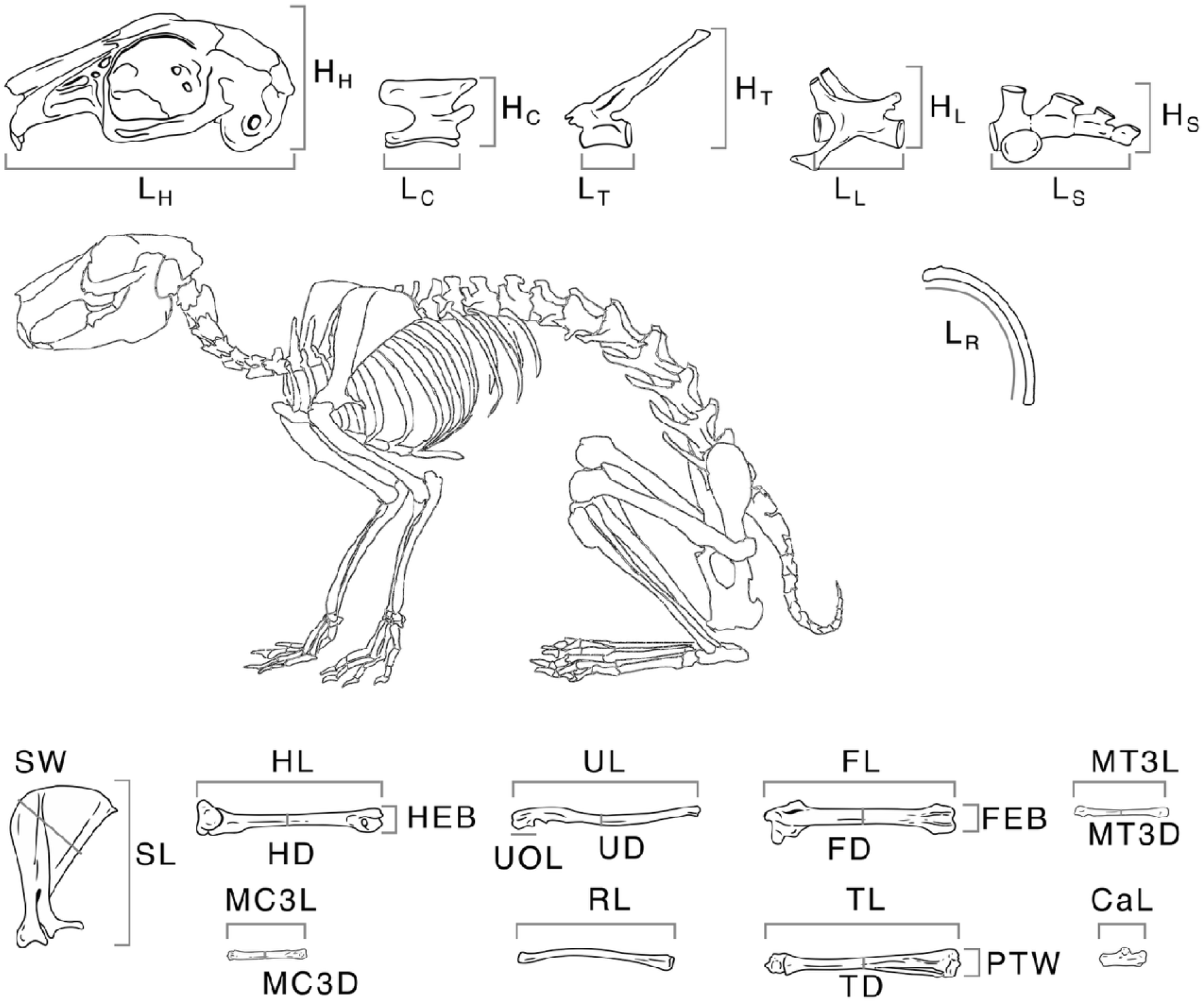
Trait measurements used to calculate head-body elongation ratio (hbER); headER; axial elongation index (AEI) of the cervical (C), thoracic (T), lumbar (L), and sacral (S) regions; and functional proxies of appendicular skeleton. See Table 1 for the list of functional proxies. Cranium and vertebrae: L_X_ = length; H_X_ = height. *Forelimb*: SL = scapula length; SW = scapula width; HL = humerus length; HD = humerus mid-shaft width; HEB = humerus distal width; UL = ulna length; UD = ulna mid-shaft width; UOL = ulnar olecranon length; RL = radius length; MC3L = third metacarpel length; MC3W = third metacarpel width. *Hindlimb*: FL = femur length; FD = femur mid-shaft width; FEB = femur distal width; TL = tibia length; TD = tibia mid-shaft width; TDW = tibia distal width; CaL = calcaneus length; MT3L = third metatarsal length; MT3D = third metatarsal width.

**Table 1.** Functional proxies capturing the functional diversity of the limbs (Davis 1964; Samuels and Van Valkenburgh 2008).

| Functional trait | Definition | Interpretation |
| --- | --- | --- |
| Forelimb |  |  |
| Scapula index (SI) | scapula length / scapula height | Describes the expansion of shoulder musculature versus contribution of scapula to limb elongation. |
| Brachial index (BI) | radius length / humerus length | Estimates relative proportions of proximal and distal elements of the forelimb. |
| Humeral robustness index (HRI) | mediolateral diameter of humerus / humerus length | Estimates robustness of the humerus and its ability to resist bending and shearing stresses. |
| Humeral epicondylar index (HEI) | epicondylar breadth of humerus / humerus length | Estimates relative area available for the origins of the forearm flexors, pronators, and supinators. |
| Olecranon length index (OLI) | olecranon process length / ulna length | Estimates relative mechanical advantage of the triceps brachii and dorsoepitrochlearis muscles used in elbow extension. |
| Ulnar robustness index (URI) | mediolateral diameter of ulna / ulna length | Estimates robustness of the ulna and its ability to resist bending and shearing stresses, and relative area available for the origin and insertion of forearm and manus flexors, pronators, and supinators. |
| Manus proportions index (MAN) | metacarpal 3 length / humerus length | Estimates relative proportions of proximal and distal elements of the forelimb, and relative size of the hand. |
| Hindlimb |  |  |
| Crural index (CI) | tibia length / femur length | Estimates relative proportions of proximal and distal elements of the hind limb. |
| Femoral robustness index (FRI) | anteroposterior diameter of femur / femur length | Estimates robustness of the femur and its ability to resist bending and shearing stresses. |
| Femoral epicondylar index (FEI) | epicondylar breadth of femur / femur length | Estimates relative area available for the origins of the gastrocnemius and soleus muscles used in extension of the knee and plantar-flexion of the pes. |
| Tibial robustness index (TRI) | mediolateral diameter of tibia / tibia length | Estimates robustness of the tibia and its ability to resist bending and shearing stresses. |
| Pes length<br>index (PES) | metatarsal 3 length /<br>femur length | Estimates relative proportions of proximal and<br>distal elements of the hind limb, and relative size of<br>the hind foot. |
| Pes length<br>index (PES) | metatarsal 3 length /<br>femur length | Estimates relative proportions of proximal and<br>distal elements of the hind limb, and relative size of<br>the hind foot. |

### Ecological and Behavioral Data

We classified the 40 lagomorph species in our sample into discrete groups based on burrowing behavior, locomotor mode, and ecotype (Fig. 2). For burrowing behavior, we created three categories–complex burrowers, simple burrowers, and non-burrowers–based on the literature and species accounts on the Animal Diversity Web (Myers et al. 2025) and the Handbook of the Mammals of the World (Wilson et al. 2016) (Table S1). Simple burrowers create short burrows, where soil is excavated to form an underground structure. Complex burrowers create extensive burrows that included underground structures, while non-burrowers do not exhibit digging behavior. For locomotor mode, we followed previous work and classified each species as either cursorial, saltatorial, or generalized (Kraatz and Sherratt 2016; Wood□Bailey et al. 2025). Cursorial species routinely run at high speed, saltatorial species display significant jumping actions, while generalist species do not exhibit these behaviors and are often described as scampering. Lastly, we classified the species using the traditional ecotype categories (i.e., hare, rabbit, and pika) as each group exhibits distinct functional morphological traits (Kraatz et al. 2021).

**Figure 2.**
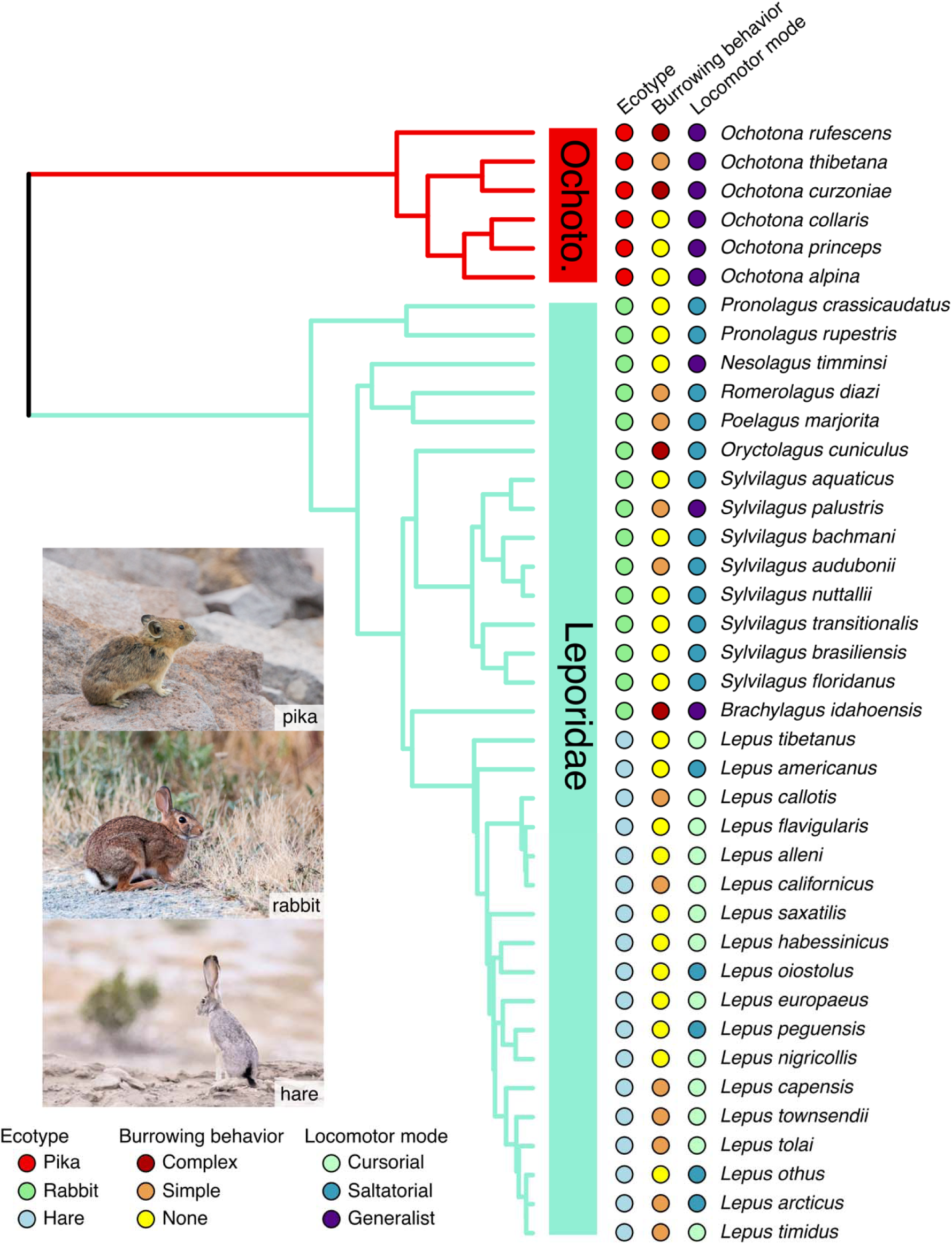
Phylogeny of lagomorph species used in this study. Photos of the pika, rabbit, and hare examples are represented by an American pika (Ochotona princeps), eastern cottontail (Sylvilagus floridanus), and black-tailed jackrabbit (Lepus californicus), respectively. All photos taken by C.J.L.

### Statistical Analyses

We conducted all analyses under a phylogenetic framework using a mammal phylogeny from (Upham et al. 2019) in R 4.3.1 (R Core Team 2023). We first determined which morphological components (i.e., cranium, ribs, cervical vertebrae, thoracic vertebrae, lumbar vertebrae, and sacrum) were most related to body shape by performing a phylogenetic multiple regression with the R package RRPP v1.4.0 (Collyer and Adams 2018). The six morphological components were used as explanatory variables and body shape was used as the response variable. We used R^2^ to examine the proportion of the variance in body shape explained by all the morphological components and determined that the component most associated with body shape had the highest R^2^. Statistical significance was determined using the random residual permutation procedure (RRPP) with 1,000 iterations (Adams and Collyer 2018). We repeated the phylogenetic multiple regression in just leporids but did not have a large enough sample size to test ochotonids.

We then examined the relationships between each limb length and body shape using phylogenetic generalized least squares (PGLS). Regression coefficients for all models were estimated simultaneously with phylogenetic signal as Pagel’s λ in the residual error using the R package phylolm v2.6.2 (Ho and Ané 2014). We also generated 1,000 bootstrap replications of the slopes in each model. We tested if the relationship between each limb and body shape was significant based on whether the 95% confidence intervals from 1,000 bootstrap replications deviated from an isometric slope of 0. We size-corrected limb lengths prior to analyses by extracting residuals for each trait against the geometric mean using a PGLS. We repeated these analyses in both leporids and ochotonids.

Lastly, we tested how size, burrowing behavior, locomotor mode, ecotype, and family influenced body shape using a series of 10 PGLS models (Table 2). Regression coefficients for all models were estimated simultaneously with phylogenetic signal as Pagel’s λ in the residual error. We evaluated our models using Akaike information criterion weights corrected for sample size (AICcW). We considered all models with ΔAICc < 2 as the best supported model(s). In instances where the best supported model(s) incorporated a discrete effect (i.e., burrowing behavior, locomotor mode, or ecotype), we assessed if body shape differed among the discrete groups by computing the observed difference between the mean values of each pair and creating its 95% confidence interval using the bootstrap replications. Confidence intervals that encompassed zero indicated that ecotype pairs were not significantly different from each other. We also generated 1000 bootstrap replications of the slopes and means of each variable in each of the best fitting models.

**Table 2.** PGLS models used to test how size, burrowing behavior, locomotor mode, ecotype, and family influenced body shape and limb functional indices.

| Model name | Model explanation |
| --- | --- |
| PGLS<sub>size</sub> | *body shape ~ ln size*: tests how each body shape scales with skeletal size |
| PGLS<sub>locomotor mode</sub> | *body shape ~ locomotory mode*: tests locomotor mode effects on body shape |
| PGLS<sub>size*locomotor mode</sub> | *body shape ~ ln size \* locomotory mode*: tests the interaction between size and locomotor mode effects on body shape |
| PGLS<sub>size+locomotor mode</sub> | *body shape ~ ln size + locomotory mode*: tests size and locomotor mode effects on body shape |
| PGLS<sub>burrowing behavior</sub> | *body shape ~ burrowing*: tests burrowing behavior effects on body shape |
| PGLS<sub>size*burrowing behavior</sub> | *body shape ~ ln size \* burrowing*: tests the interaction between size and burrowing behavior effects on body shape |
| PGLS<sub>size+burrowing behavior</sub> | *body shape ~ ln size + burrowing*: tests size and burrowing behavior effects on body shape |
| PGLS<sub>ecotype</sub> | *body shape ~ ecotype*: tests ecotype effects on body shape |
| PGLS<sub>size*ecotype</sub> | *body shape ~ ln size \* ecotype*: tests the interaction between size and ecotype effects on body shape |
| PGLS<sub>size+ecotype</sub> | *body shape ~ ln size + ecotype*: tests size and ecotype effects on body shape |

We repeated this model selection procedure for each body shape component (i.e., headER; relative rib length; and cervical, thoracic, lumber, and sacral AEI) and each functional proxy of the limbs.

## Results

### Morphological components underlying body shape

Across lagomorphs, we found that body shape was best predicted by relative length of the ribs (R^2^ = 0.73; P = 0.001) and elongation or shortening of the thoracic (R^2^ = 0.11; P = 0.001) and lumbar (R^2^ = 0.02; P = 0.015) regions (Fig. 3; Table 3). Elongation or shortening of the head, cervical, and sacral regions combined explained less than 2%.

**Fig. 3.**
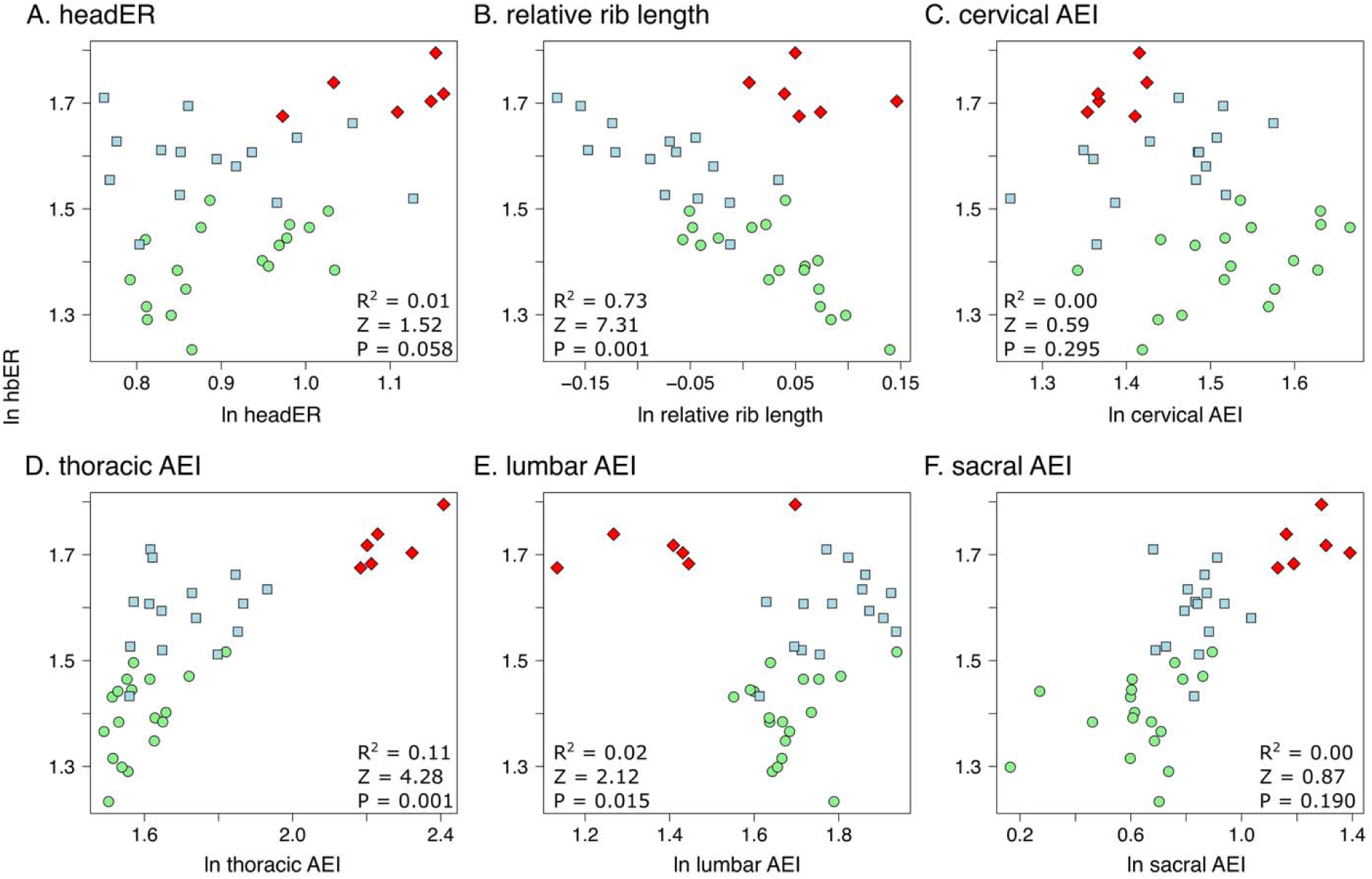
Scatterplots of head-body elongation ratio (hbER) versus skeletal components underlying body shape diversity. R^2^ and Z scores were obtained from phylogenetic multiple regression with the random residual permutation procedure (RRPP). See Table 3 for full results. Colors are the same as in Figure 3.

**Table 3.** Results of the phylogenetic multiple regression to determine which morphological components contributed most to body shape evolution across all lagomorphs. Bold p-values indicate significance (α = 0.05). DF, degrees of freedom; SS, sum of squares.

| Clade | Body shape component | Df | SS | MS | Rsqr | F | Z | P |
| --- | --- | --- | --- | --- | --- | --- | --- | --- |
| All lagomorphs |  |  |  |  |  |  |  |  |
|  | In head elongation ratio | 1 | 0.000 | 0.000 | 0.01 | 3.51 | 1.52 | 0.058 |
|  | **In relative rib length** | **1** | **0.040** | **0.040** | **0.73** | **322.19** | **7.31** | **0.001** |
|  | In cervical AEI | 1 | 0.000 | 0.000 | 0.00 | 1.28 | 0.59 | 0.295 |
|  | **In thoracic AEI** | **1** | **0.006** | **0.006** | **0.11** | **46.41** | **4.28** | **0.001** |
|  | **In lumbar AEI** | **1** | **0.001** | **0.001** | **0.02** | **7.49** | **2.12** | **0.015** |
|  | In sacral AEI | 1 | 0.000 | 0.000 | 0.00 | 1.83 | 0.87 | 0.190 |
|  | Residuals | 32 | 0.004 | 0.000 | 0.07 |  |  |  |
|  | Total | 38 | 0.055 |  |  |  |  |  |
| Leporidae |  |  |  |  |  |  |  |  |
|  | In head elongation ratio | 1 | 0.000 | 0.000 | 0.01 | 2.48 | 1.15 | 0.137 |
|  | **In relative rib length** | **1** | **0.037** | **0.037** | **0.69** | **186.14** | **6.46** | **0.001** |
|  | In cervical AEI | 1 | 0.000 | 0.000 | 0.00 | 0.36 | -0.11 | 0.563 |
|  | **In thoracic AEI** | **1** | **0.006** | **0.006** | **0.12** | **32.75** | **3.71** | **0.001** |
|  | **In lumbar AEI** | **1** | **0.002** | **0.002** | **0.03** | **8.95** | **2.46** | **0.004** |
|  | **In sacral AEI** | **1** | **0.001** | **0.001** | **0.02** | **5.05** | **1.66** | **0.037** |
|  | Residuals | 26 | 0.005 | 0.000 | 0.10 |  |  |  |
|  | Total | 32 | 0.053 |  |  |  |  |  |
| Ochotonidae |  |  |  |  |  |  |  |  |
| *Not a large enough sample size to test* |  |  |  |  |  |  |  |  |

Leporids exhibited similar trends. Leporid body shape was best predicted by relative length of the ribs (R^2^ = 0.69; P = 0.001) and elongation or shortening of the thoracic (R^2^ = 0.12; P = 0.001), lumbar (R^2^ = 0.03; P = 0.004), and sacral (R^2^ = 0.02; P = 0.037) regions. We did not have a large enough sample size to test ochotonids.

### Relationships between body shape and limb length

We found negative trends between lagomorph body shape and relative limb lengths, in which lagomorphs with more elongate bodies tend to exhibit relatively shorter forelimbs and hindlimbs (Fig. 4). However, the slopes of these relationships were not statistically significantly different from 0 in either size-corrected forelimb length (R^2^ = 0.10, λ = 0.00, P = 0.054; slope [95% CI] = −0.15 [−0.29:0.01]) or size-corrected hind limb length (R^2^ = 0.01, λ = 0.00, P = 0.621; slope = −0.05 [−0.23:0.12]; Fig. 4).

**Fig. 4.**
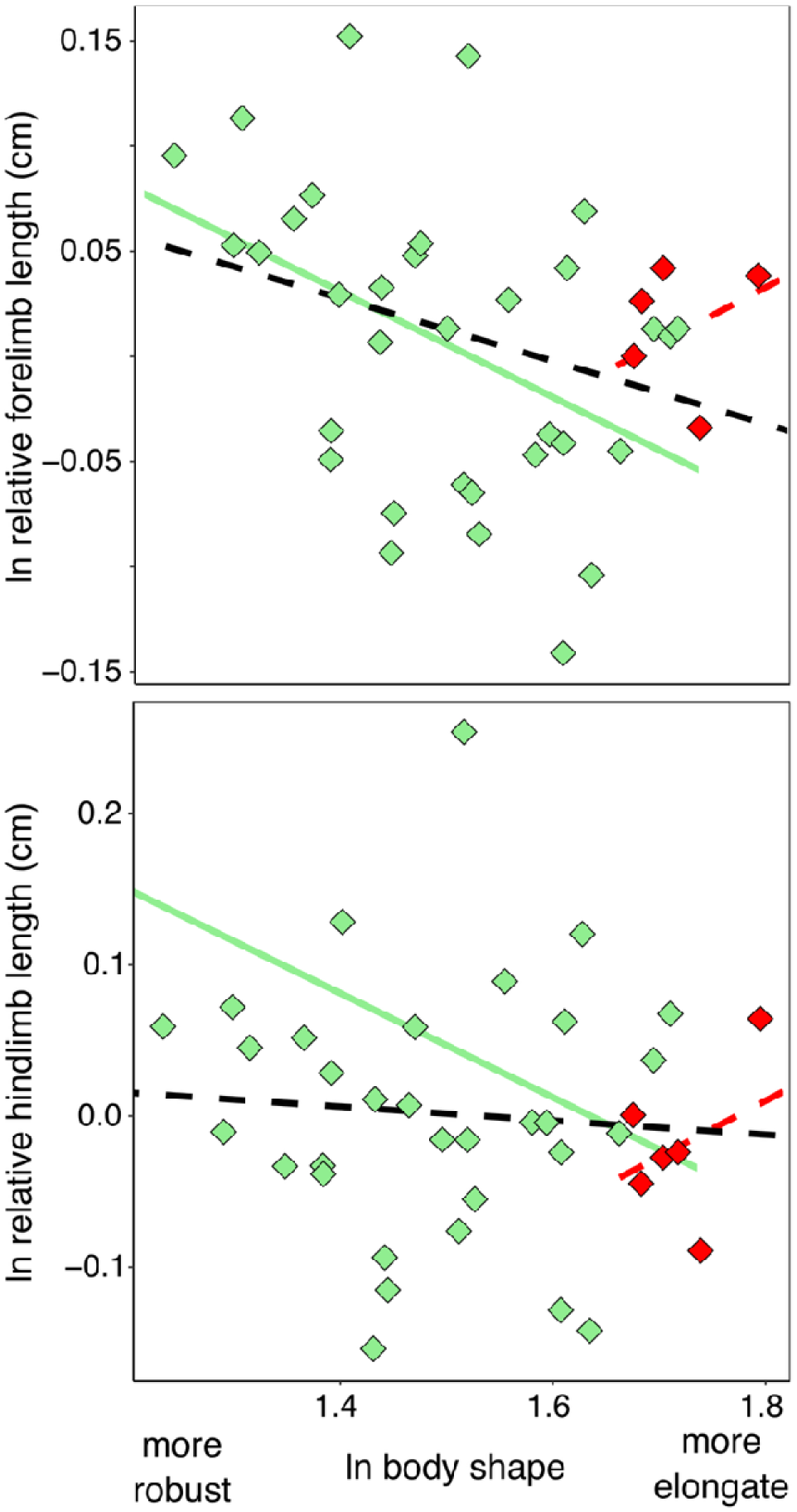
Plots of PGLS regressions showing the relationship between ln body shape and ln relative (i.e., size-corrected) forelimb and hind limb lengths across lagomorphs (black line) and within leporids (green line) and ochotonids (red line). Dashed lines indicate that confidence intervals generated by 1,000 bootstrap replications indicated that slopes did not significantly differ from 0.

Leporids also exhibited negative trends between body shape and relative limb lengths, where leporids with more elongate bodies exhibited relatively shorter forelimbs (R^2^ = 0.18, λ = 0.00, P = 0.013; slope = −0.25 [−0.44:-0.06]) and hindlimbs (R^2^ = 0.18, λ = 0.00, P = 0.013; slope = −0.25 [−0.45:-0.06]). Ochotonids trended towards positive relationships between body shape and relative limb lengths; however, the slopes of these relationships were not statistically significantly different from 0 in either size-corrected forelimb length (R^2^ = 0.11, λ = 0.00, P = 0.251; slope = 0.15 [−0.03:0.33]) or size-corrected hind limb length (R^2^ = 0.31, λ = 0.76, P = 0.251; slope = 0.15 [−0.05:0.31]).

### Predictors of body shape and underlying components

The PGLS_size_ model was best supported for hbER (R^2^ = 0.34, AICcW = 0.95; Table S2), indicating that larger lagomorphs exhibited more robust bodies whereas smaller lagomorphs exhibited more elongate bodies (Fig. 5).

**Fig. 5.**
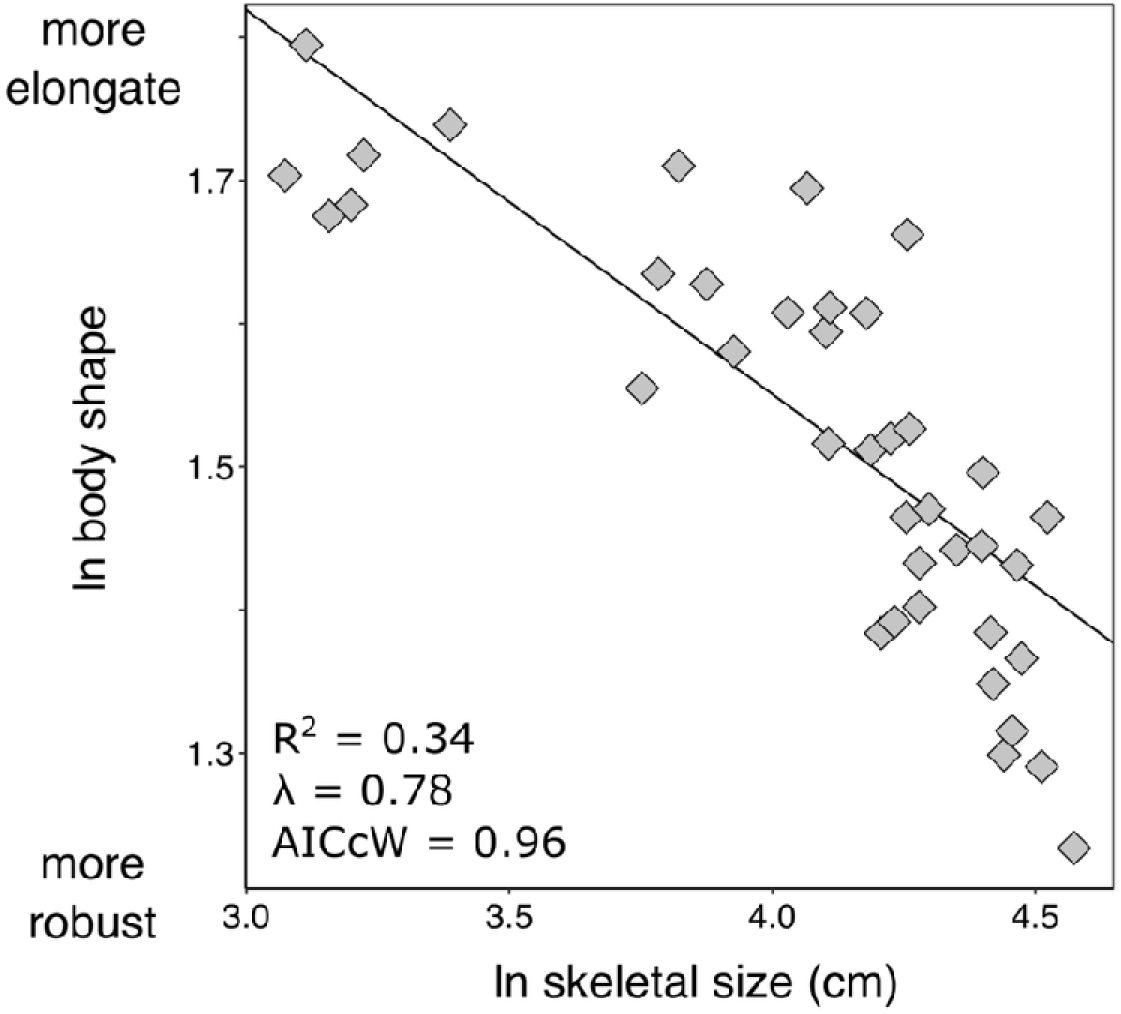
Graphical representation of the best supported model (PGLS_size_ model) for lagomorph body shape diversity. See Table 4 for full parameter estimates.

**Table 4.**
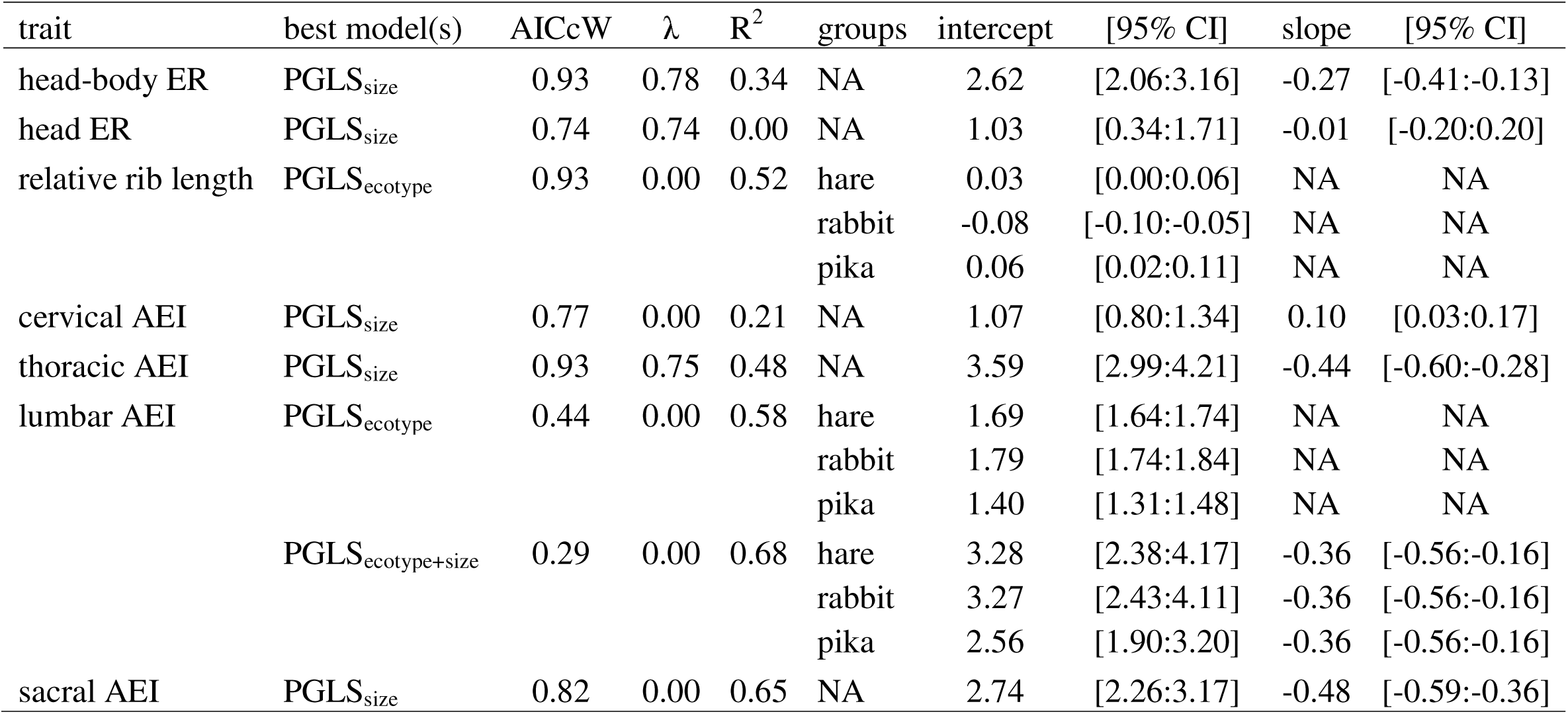
Parameter estimates from best supported model(s) for body shape and underlying skeletal components. 95% confidence intervals for intercepts and slopes (if applicable) were generated using 1000 bootstrap replications.

Within the body shape components, the PGLS_size_ model was also the best supported model for head elongation (R^2^ = 0.00, AICcW = 0.74), cervical AEI (R^2^ = 0.21, AICcW = 0.77), thoracic AEI (R^2^ = 0.48, AICcW = 1.00), and sacral AEI (R^2^ = 0.65, AICcW = 0.7782 Fig. 6; Table S2). Larger lagomorphs exhibited more elongate cervical regions (i.e., higher cervical AEI) but more robust thoracic and sacral regions (i.e., lower thoracic and sacral AEI; Fig. 6; Table 4). Larger lagomorphs also trended towards more robust heads, but this relationship was not significant (Fig. 6; Table 4). Relative rib length was best supported by the PGLS_ecotype_ model (R^2^ = 0.52, AICcW = 0.93; Fig. 6; Table S2). Rabbits exhibited significantly shorter relative rib length (mean rib [95% confidence interval] = −0.08 [−0.10:-0.05]) compared to hares (0.03 [0.01:0.06]) and pikas (0.06 [0.02:0.10]) (Table 4; Table S3). We found equal support for the PGLS_ecotype_ (R^2^ = 0.58, AICcW = 0.44) and PGLS_ecotype+size_ (R^2^ = 0.68, AICcW = 0.29) models for lumbar AEI. All three ecotypes exhibited more robust lumbar regions with increasing size, but pikas (1.40 [1.31:1.48]) exhibited relatively more robust lumbar regions than hares (1.69 [1.64:1.74]) and rabbits (1.79 [1.74:1.84]) (Fig. 6; Table 4; Table S3).

**Fig. 6.**
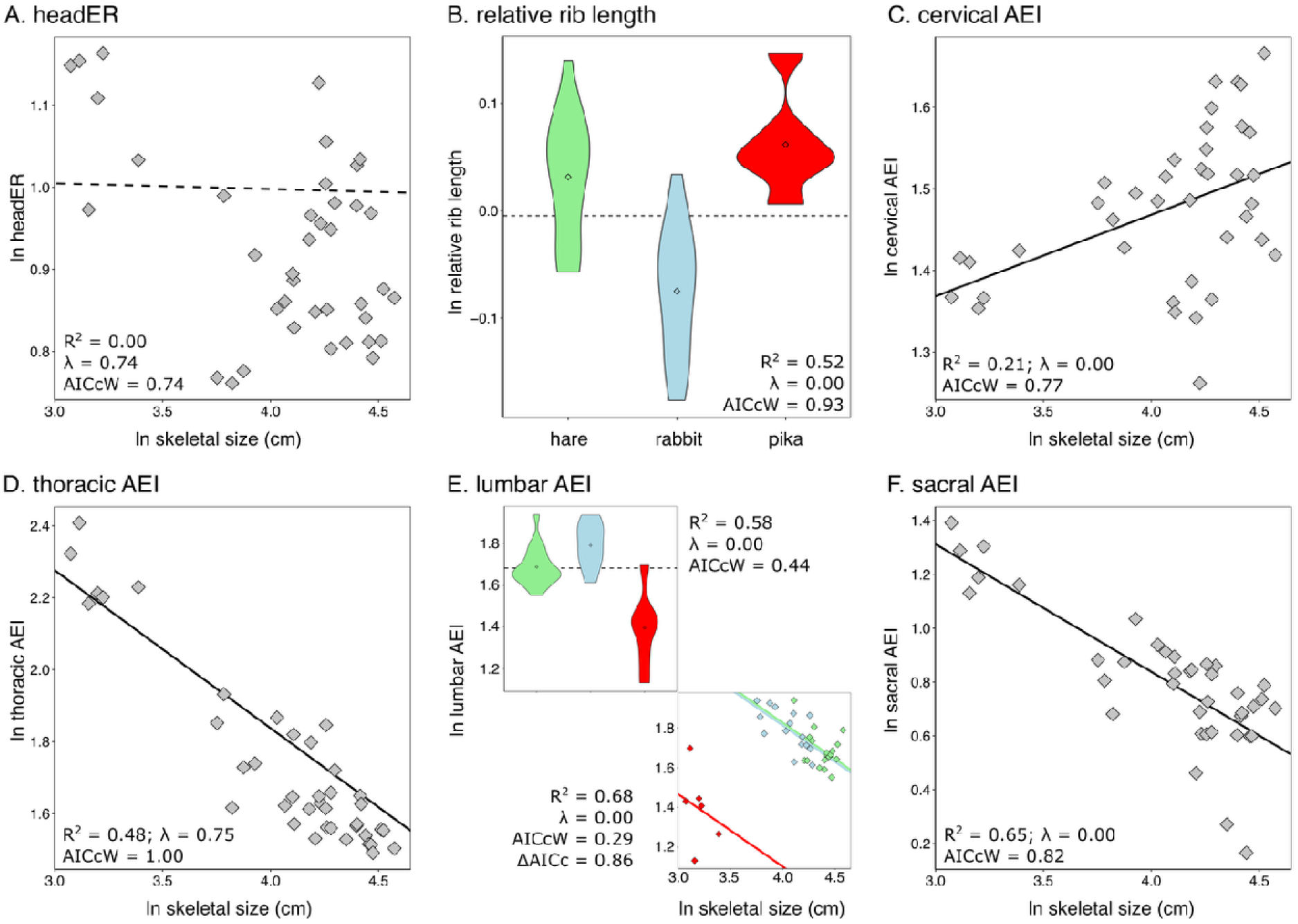
Graphical representations of the best supported model(s) for skeletal components underlying body shape. Colors are the same as in Figure 2. See Table 4 for full parameter estimates.

### Predictors of limb functional proxies

We found equal support for the PGLS_size_ and PGLS_ecotype_ models in the BI, OLI, FEI, and PES indices (Fig. 7, 8; Table S4). Under the PGLS_ecotype_ model, pikas exhibited the lowest BI (0.78 [0.72:0.83]) and PES (0.37 [0.35:0.40]) indices, followed by rabbits (BI = 0.91 [0.88:0.95]; PES = 0.41 [0.40:0.43]), then hares (BI = 1.09 [1.06:1.12]; PES= 0.45 [0.44:0.47]) (Fig. 7; Table 5; Table S5). Consistently, larger lagomorphs trended towards increased BI and PES under the PGLS_size_ model, but these relationships were not statistically significant (Fig. 7; Table 5). In contrast, smaller lagomorphs exhibited more robust femoral epicondyle, and pikas exhibited the highest OLI (pika = 0.13 [0.13:0.14], rabbit = 0.11 [0.11:0.12], hare = 0.09 [0.09:0.10]) (Table 5; Table S5). (Fig. 7; Table 5).

**Fig. 7.**
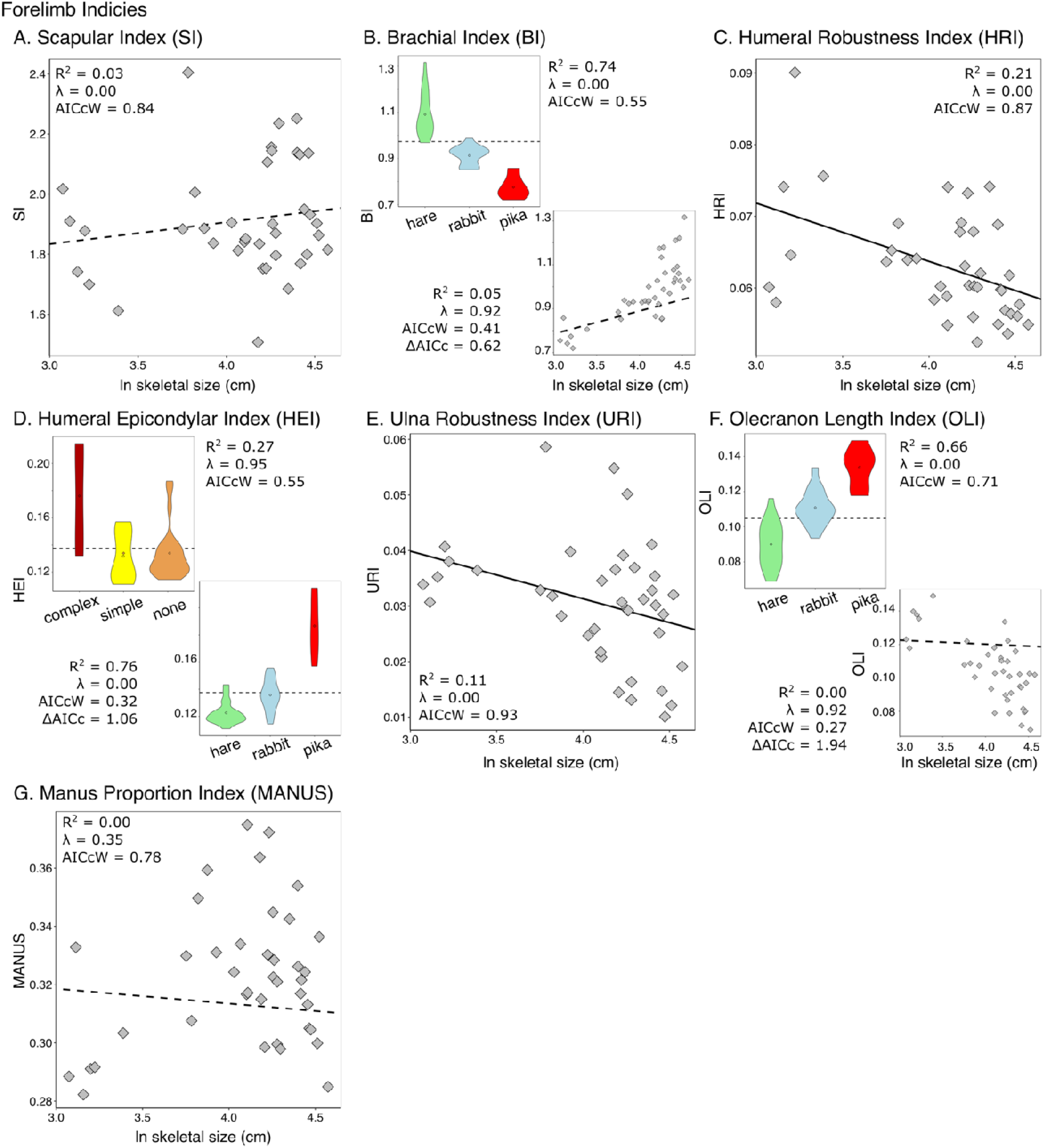
Graphical representations of the best supported model(s) for functional proxies of the forelimb. Colors are the same as in Figure 2. See Table S4 for model comparisons and Table 5 for full parameter estimates.

**Table 5.** Parameter estimates from best supported models for limb functional proxies. 95% confidence intervals for intercepts and slopes (if applicable) were generated using 1000 bootstrap replications.

| Limb indices | best model(s) | AICcW | $\lambda$ | R<sup>2</sup> | groups | intercept | [95% CI] | slope | [95% CI] |
| --- | --- | --- | --- | --- | --- | --- | --- | --- | --- |
| Scapula index (SI) | PGLS<sub>size</sub> | 0.84 | 0.00 | 0.03 | NA | 1.62 | [1.00 : 2.20] | 0.073 | [-0.073 : 0.217] |
| Brachial index (BI) | PGLS<sub>ecotype</sub> | 0.55 | 0.00 | 0.74 | hare | 1.09 | [1.06 : 1.12] | NA | NA |
|  |  |  |  |  | rabbit | 0.91 | [0.88 : 0.95] | NA | NA |
|  |  |  |  |  | pika | 0.78 | [0.72 : 0.84] | NA | NA |
|  | PGLS<sub>size</sub> | 0.41 | 0.92 | 0.1 | NA | 0.47 | [-0.11 : 1.15] | 0.104 | [-0.067 : 0.256] |
| Humeral Robustness Index (HRI) | PGLS<sub>size</sub> | 0.87 | 0.00 | 0.21 | NA | 0.10 | [0.08 : 0.12] | -0.008 | [-0.014 : -0.003] |
| Humeral Epicondylar Index (HEI) | PGLS<sub>burrow</sub> | 0.55 | 0.95 | 0.27 | complex | 0.18 | [0.15 : 0.21] | NA | NA |
|  |  |  |  |  | simple | 0.15 | [0.12 : 0.18] | NA | NA |
|  |  |  |  |  | none | 0.15 | [0.12 : 0.18] | NA | NA |
|  | PGLS<sub>ecotype</sub> | 0.32 | 0.00 | 0.76 | hare | 0.12 | [0.12 : 0.13] | NA | NA |
|  |  |  |  |  | rabbit | 0.14 | [0.13 : 0.14] | NA | NA |
|  |  |  |  |  | pika | 0.19 | [0.18 : 0.20] | NA | NA |
| Olecranon Length Index (OLI) | PGLS<sub>ecotype</sub> | 0.71 | 0.00 | 0.66 | hare | 0.09 | [0.09 : 0.10] | NA | NA |
|  |  |  |  |  | rabbit | 0.11 | [0.11 : 0.12] | NA | NA |
|  |  |  |  |  | pika | 0.13 | [0.12 : 0.14] | NA | NA |
|  | PGLS<sub>size</sub> | 0.27 | 0.92 | 0.00 | NA | 0.13 | [0.04 : 0.23] | -0.002 | [-0.030 : 0.023] |
| Ulna Robustness Index (URI) | PGLS<sub>size</sub> | 0.93 | 0.00 | 0.11 | NA | 0.07 | [0.03 : 0.09] | -0.009 | [-0.017 : -0.001] |
| Manus Proportion Index (MAN) | PGLS<sub>size</sub> | 0.78 | 0.35 | 0.00 | NA | 0.33 | [0.21 : 0.45] | -0.005 | [-0.036 : 0.029] |
| Crural Index (CI) | PGLS<sub>ecotype</sub> | 0.69 | 0.00 | 0.3 | hare | 1.19 | [1.17 : 1.22] | NA | NA |
|  |  |  |  |  | rabbit | 1.11 | [1.08 : 1.13] | NA | NA |
|  |  |  |  |  | pika | 1.15 | [1.11 : 1.19] | NA | NA |
| Femoral Robustness Index (FRI) | PGLS<sub>size</sub> | 0.81 | 0.00 | 0.1 |  | 0.10 | [0.07 : 0.12] | -0.004 | [-0.009 : 0.001] |
| Femoral Epicondylar Index (FEI) | PGLS<sub>size</sub> | 0.69 | 0.00 | 0.3 | NA | 0.24 | [0.20 : 0.28] | -0.018 | [-0.029 : -0.008] |
|  | PGLS<sub>ecotype</sub> | 0.39 | 0.00 | 0.30 | hare | 0.16 | [0.16 : 0.17] | NA | NA |
|  |  |  |  |  | rabbit | 0.17 | [0.16 : 0.17] |  |  |
|  |  |  |  |  | pika | 0.19 | [0.18 : 0.20] | NA | NA |
| Tibial Robustness Index (TRI) | PGLS<sub>size</sub> | 0.89 | 0.92 | 0 | NA | 0.02 | [-0.06 : 0.09] | 0.012 | [-0.007 : 0.032] |
| Pes Proportion Index (PES) | PGLS<sub>ecotype</sub> | 0.62 | 0.00 | 0.7 | hare | 0.45 | [0.44 : 0.47] | NA | NA |
|  |  |  |  |  | rabbit | 0.41 | [0.39 : 0.43] | NA | NA |
|  |  |  |  |  | pika | 0.37 | [0.35 : 0.40] | NA | NA |
|  | PGLS<sub>size</sub> | 0.36 | 0.57 | 0.00 | NA | 0.35 | [0.11 : 0.57] | 0.013 | [-0.045 : 0.075] |
| Intermembral Index (IM) | PGLS<sub>size</sub> | 0.81 | 0.88 | 0.2 | NA | 0.31 | [-0.03 : 0.65] | 0.111 | [0.030 : 0.203] |

The PGLS_size_ model was the best supported model of the SI, MANUS, IM, and limb robustness indices (Fig. 7, 8; Table S4). Larger lagomorphs exhibited more gracile humeri and ulna (Fig. 7) and trended towards more gracile femurs but more robust tibia; however, these relationships in the hindlimb were not statistically significant (Fig. 8; Table 5). Similarly, larger lagomorphs also trended towards more robust scapula and relatively shorter metacarpals, but these relationships were not statistically significant (Fig. 7; Table 5). The PGLS_ecotype_ model was the best supported model for the CI index (Fig. 7; Table S4). Hares (1.19 [1.17:1.21]) and rabbits (1.11 [1.08:1.13]) exhibited significantly different CI but pikas (1.15 [1.11:1.19]) did not differ from either group (Fig. 7; Table 5; Table S5).

**Fig. 8.**
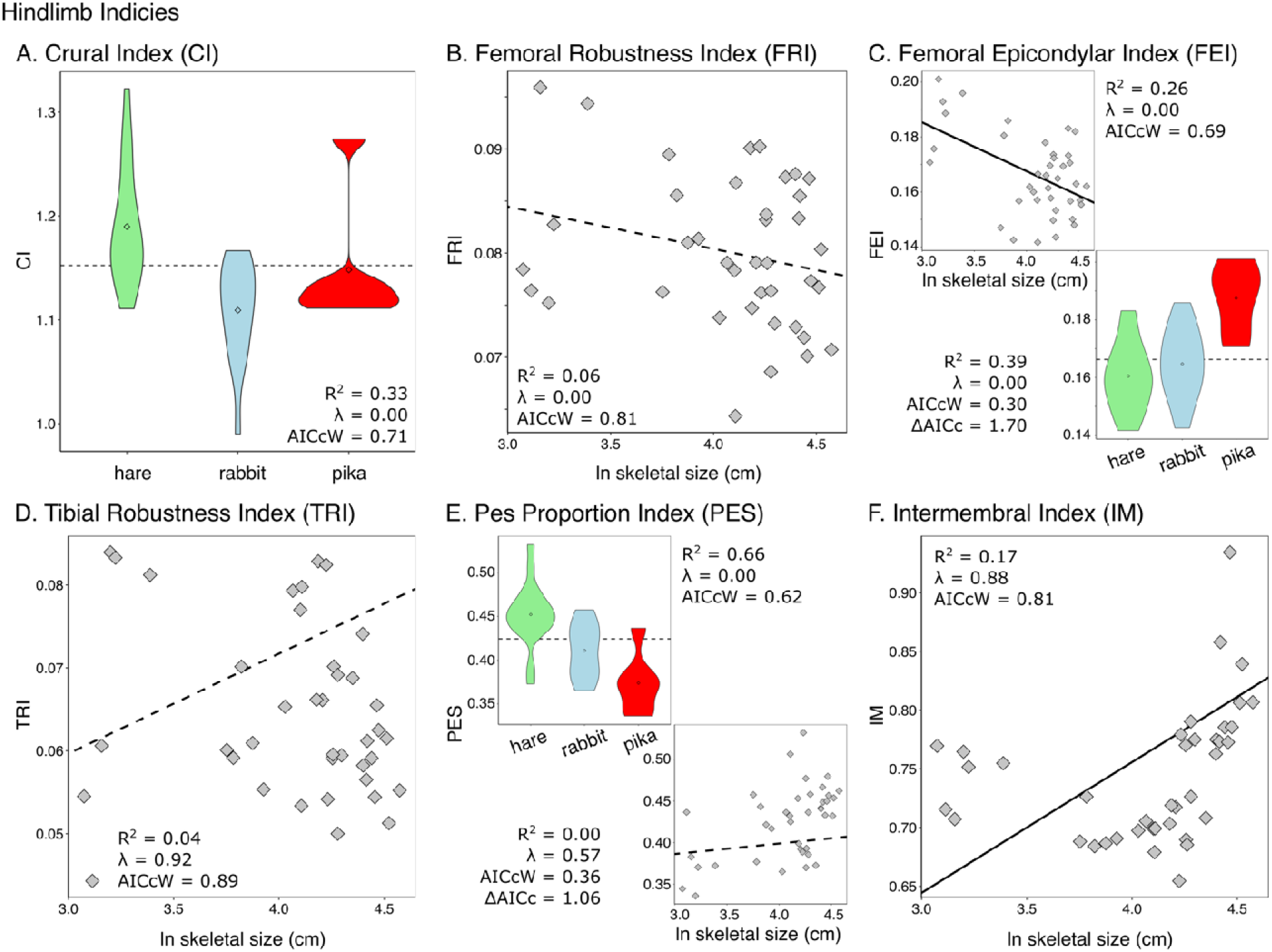
Graphical representations of the best supported model(s) for functional proxies of the hindlimb. Colors are the same as in Figure 2. See Table S4 for model comparisons and Table 5 for full parameter estimates.

Lastly, we found equal support for the PGLS_burrow_ and PGLS_ecotype_ models for the HEI index (Fig. 7B; Table S4). Lagomorphs with complex burrows exhibited significantly higher HEI (0.18 [0.15:2.13]) than lagomorphs with simple burrows (0.15 [0.12:0.18]) or that do not burrow (0.15 [0.12:0.18]), and there was no difference in HEI between simple burrowers and non-burrowers (Fig. 7; Table 5; Table S5). Under the PGLS_ecotype_ model, pikas exhibited the highest HEI 0.19 [0.15:0.20], followed by rabbits (0.14 [0.13:0.14]) and hares (0.12 [0.12:0.13]) (Fig. 7; Table 5; Table S5).

## Discussion

### Components underlying body shape and limb length evolution

The patterns underlying body shape diversity are poorly documented in mammals. This is in contrast to ectothermic vertebrates, in which a plethora of studies have revealed the multiple pathways in which the cranial and axial components contribute to body shape evolution as well as associated evolutionary scaling of the appendicular skeleton with changes in body shape (Gans 1975; Parra-Olea and Wake 2001; Wiens and Slingluff 2001; Ward and Brainerd 2007; Skinner et al. 2008; Ward and Mehta 2010; Collar et al. 2013). Investigation of the skeletal components underlying mammalian body shape variation and associated evolutionary changes in limb lengths has only been conducted in carnivorans and squirrels (Law et al. 2019; Law 2021b; Linden et al. 2023). These studies found that shortening/elongation of the lumbar region (R^2^ = 0.41) followed by relative depth of the ribcage (R^2^ = 0.21) and shortening/elongation of the thoracic region (R^2^ = 0.14) contributed the most to body shape variation in carnivorans (Law 2021b) whereas only relative depth of the ribcage (R^2^ = 0.21) and shortening/elongation of the thoracic region (R^2^ = 0.14) contributed the most to body shape variation in squirrels (Linden et al. 2023). In this study, we found that relative depth of the ribcage (R^2^ = 0.73) followed by shortening/elongation of the thoracic (R^2^ = 0.11) and lumbar (R^2^ = 0.02) regions were significant predictors of lagomorph body shape variation, contributing 86% of total lagomorph body shape variation. Thus, in all three clades, evolutionary changes in body shapes were primarily achieved through elongation/shortening of the thoracolumbar region. This may be unsurprising considering that the thoracolumbar region of the vertebral column and ribcage are the primary structures used to support the body against gravity and transmit and receive propulsive forces to/from the limbs for locomotion (Kardong 2014). Elongation or shortening of the lumbar region may also facilitate maneuverability or stability, respectively, of the posterior vertebral column, which in turn may contribute to distinct locomotor behaviors (Boszczyk et al. 2001; Jones 2015b,a). Evolutionary changes in the lumbar region were important contributors to body shape variation in carnivorans (and to a lesser extent in lagomorphs) and may suggest that cursoriality and other locomotor behaviors requiring vertebral mobility influences body shape evolution. However, (Law 2021a) found weak support that locomotor mode influenced evolutionary changes in the lumbar region, attributing this lack of relationship to the fact that the AEI metric is a simple ratio between vertebral length and height and cannot capture finer morphological features of the vertebral column. Alas, ecomorphological analyses revealed the evolution of specific vertebral traits and overall vertebral shape are influenced by different locomotor modes and hunting behaviors across carnivorans (Figueirido et al. 2021; Law et al. 2024). A similar scenario probably also explains lagomorphs; elongation ratio metrics such as hbER and AEI does not capture the mobility or stability of vertebral joints (see next section). Thus, future work is necessary to test relationships among overall body shape, the underlying skeletal components, and other metrics that capture vertebral joints, shape, and material properties in lagomorphs.

In investigations of relationships between body shape and limb lengths, researchers have revealed that more elongate species tend to exhibit relatively shorter or even a loss of limbs/fins (Gans 1975; Parra-Olea and Wake 2001; Wiens and Slingluff 2001; Ward and Brainerd 2007; Skinner et al. 2008; Ward and Mehta 2010). The few studies that have examined this relationship in mammals found similar patterns in the forelimbs but not hindlimbs of musteloids and ground squirrels; specifically, more elongate musteloids and ground squirrels exhibited relatively shorter forelimbs but not hindlimbs and are smaller bodied (Law 2019; Linden et al. 2023). Here, we found a significant relationship between more elongate body shapes and relatively shorter forelimbs and hindlimbs in leporids (Fig. 4). However, these patterns were not significant in ochotonids as well as across all lagomorphs. The combination of smaller, more elongate bodies with relatively shorter forelimbs is often associated with the evolution of innovative locomotor and foraging behaviors (Webb 1982; Gans 1983; Brainerd and Patek 1998; Bergmann and Irschick 2009; Mehta et al. 2010). In mammals, these traits are hypothesized to enable weasels to chase prey through winding underground burrows or tight crevices and for ground squirrels to dig large burrow systems (Law 2019; Linden et al. 2023). However, most leporids do not utilize or dig complex burrows to the extent that weasels or ground squirrels do. Therefore, perhaps robust leporids rather than elongate species are dictating the relationship between body shape and relative limb length in this clade. Robust leporids tend to be large, cursorial hares (Fig. 1; Fig. 5), and having relatively longer limbs would lead to reduced energetic cost of transport and increased running speed by increasing stride length and decreasing the moment of inertia of the limbs (Kram and Taylor 1990; Strang and Steudel 1990; Garland and Janis 1993; Polly 2007; Pontzer 2007b,a; Kilbourne and Hoffman 2013). Correspondingly, we found that the largest lagomorphs display several adaptations in their limbs to facilitate cursoriality (see next section below).

### The evolution of lagomorph body shape and appendicular skeleton

Evolutionary changes in body size have a strong influence on variation of morphological traits (Bertram and Biewener 1990; Marroig and Cheverud 2005; Martín-Serra et al. 2014; Law 2021b; Holding et al. 2022; Mitchell et al. 2023). Consistently, we found that PGLS models incorporating body size were the best supported models explaining variation in body shape and many limb functional proxies in lagomorphs. Specifically, larger lagomorphs tend to exhibit more robust body shapes but more gracile forelimbs, whereas smaller lagomorphs tend to exhibit more elongate body shapes but more robust forelimbs. These results may be surprising considering that locomotion via the cursoriality gradient is often hypothesized to influence skeletal variation in lagomorphs (Camp and Borell 1937; Young et al. 2014; Martin et al. 2022). The likely reason is that previous analyses of lagomorph ecomorphology tend to remove the effects of size using residuals or log-shape ratios to focus on relationships between size-corrected traits and ecological and/or behavioral factors. Thus, how scaling influences variation in lagomorph skeletal traits in addition to ecological or behavioral factors were rarely assessed in lagomorphs.

Body size was the best predictor of shape variation in the overall body, and thoracic and lumbar regions (Fig. 5, 6; Table S2). We found that robusticity of the body as well as the thoracic and lumbar regions increased with increasing size, following a trend found in other mammals such as terrestrial carnivorans and ground squirrels (Law 2021a; Linden et al. 2023). Robust body shapes along with more robust thoracolumbar centra provides increased support of the axial skeleton for larger mammals and their heavier bodies (Halpert et al. 1987; Kardong 2014; Jones 2015a,b), and our results suggest that lagomorphs are not an exception. However, the largest lagomorphs (i.e., hares) tend to exhibit cursorial locomotion; thus, it is somewhat surprising that these lagomorphs do not exhibit more elongate bodies and elongation of the thoracolumbar region, adaptations typically thought to lead to increased dorsoventral flexibility and maneuverability for greater cursorial efficiency (Boszczyk et al. 2001; Jones 2015a,b; Law et al. 2019; Linden et al. 2023). The likely explanation is that our body shape metric does not capture the sagittal mobility of the vertebral joints in cursorial lagomorphs. Lagomorph lumbar vertebrae exhibit high range of motion in the sagittal plane and low axial twisting, which facilitates extensive dorsoventral flexion and extension of the body axis (Grauer et al. 2000). High sagittal flexion and extension are hypothesized to contribute to the diversity of mammalian locomotion, particularly asymmetrical gaits such as gallop, half-bound, and bound (Hildebrand 1959; Gambaryan 1974; Hildebrand 1985b; Schilling and Hackert 2006). As half-bounding mammals, the lagomorph lumbar region is essential for locomotion, where extension and flexion of the lumbar region is paired with hindlimb thrust and hindlimb protracting simultaneously with forelimb landing, respectively (Gambaryan 1974). Thus, further investigation of the vertebral joints across lagomorphs may provide a better understanding of how backbone flexibility can be achieved despite having relatively robust bodies.

Proxies of forelimb robustness were also best predicted by size, where humeral and ulnar robustness decreased with increasing size (Fig. 7C, E; Table S4). Across Mammalia, larger species tend to exhibit more robust limb bones to accommodate the disproportionate increase in body mass relative to bone surface area at larger sizes (Biewener 1983; Bertram and Biewener 1990; Christiansen 1999; Campione and Evans 2012). However, lagomorphs diverged from this general trend, exhibiting increasingly more gracile humeri and ulnae with increased body size (Fig 7C, E; Table S4). This distinctive pattern likely reflects cursoriality in the largest lagomorphs. Long, gracile limb bones facilitate increased stride length and decreased moment of inertia of limbs, in turn decreasing the energetic cost of transport and increasing running speed (Kram and Taylor 1990; Strang and Steudel 1990; Garland and Janis 1993; Polly 2007; Pontzer 2007b,a; Kilbourne and Hoffman 2013). Furthermore, the largest lagomorphs, particularly hares, exhibited relatively longer radii compared to the humerus (high brachial index (BI); Fig. 7B; (Martin et al. 2022). Relative lengthening of the distal limb bones also increases stride length for better running efficiency. Thus, relatively long, gracile forelimbs, particularly at the distal ends, may help contribute to fast running speeds in these large lagomorphs. Nevertheless, that the PGLS models incorporating locomotor mode were not as well supported for the forelimb robusticity and brachial indices (ΔAICc = 5.92–44.76; Table S4) suggest that there is notable variation in these indices within cursorial, saltatorial, and generalized lagomorphs.

The largest lagomorphs, particularly hares, also exhibited longer tibia and third metatarsal relative to femoral length (Fig. 8A, E; Table 5, S5), suggesting similar contributions of the hindlimb towards increasing stride length. In turn, the largest lagomorphs also exhibited intermembral indices closest to 1.0, indicating nearly equal lengths of the forelimbs and hindlimbs (Fig. 8F; Table S4). However, femoral and tibial robustness does not decrease with increasing body size as found in the forelimb (Fig. 8B, D; Table S4). A possible explanation is that cursorial lagomorphs exhibit an evolutionary tug-of-war in evolving gracile hindlimbs for efficient cursoriality but not too gracile to resist fracture and support their larger body sizes. Because propulsion for lagomorph cursoriality is primarily generated by the hindlimbs, there may be a limit to how gracile the hindlimb bones can evolve to avoid fracture risks as relatively more robust limb bones provide resistance to bending and shearing stresses. Furthermore, the risk of fracture also increases in larger, heavier bodies (Kram and Taylor 1990; Strang and Steudel 1990; Garland and Janis 1993; Polly 2007; Pontzer 2007b,a; Kilbourne and Hoffman 2013), and as the largest lagomorphs, cursorial species may exhibit more stress on their limb bones while running, leading to potential fracture. Thus, their heavier bodies may also limit the gracileness of their hindlimb bones. Another possible adaptation to mitigate the risk of bone fracture is increased mechanical advantage of the hindlimb joints. More force-modified joints may reduce fracture risks by reducing the stress placed on the bone without the expense of reducing muscle force (Biewener 1989). *Lepus californicus* and *L. europaeus* (as representatives of cursorial lagomorphs) exhibit higher mechanical advantage of these joints than expected, a pattern opposing their prediction that the joints would be velocity-modified (low mechanical advantage) for sustained joint mobility of cursoriality (Smith and Savage 1956; Young et al. 2014; Martin et al. 2022). Increased mechanical advantage of the hindlimb joints and body sizes are also associated with a shift from a crouched to an upright posture to reduce maximum loading during locomotion (Biewener 1989). Although lagomorphs display a crouched posture, (Martin et al. 2022) noted that large lagomorphs fall within the range (3-4 kg) where limb kinematics and posture begin to shift (Fischer et al. 2002; Schmidt and Fischer 2009). Thus, how the unique skeletal features found in lagomorphs enables them to maintain crouched postures compared to other similarly sized mammals requires further investigation. Furthermore, in addition to competition from ungulate-type herbivores (Tomiya and Miller 2021), the skeletal morphology possibly also limits cursorial lagomorphs from evolving larger body sizes found in other herbivores.

Burrowing behavior exhibited limited effects on lagomorph body shape and limb ecomorphology. The humeral epicondylar index (HEI) was the only index to be best supported by a burrowing model. Under the PGLS_burrowing behavior_ model, complex burrowing lagomorphs exhibited the highest HEI values whereas simple burrowing lagomorphs and non-burrowing lagomorphs tended to exhibit similar HEI values (Fig. 7D; Table 5, S5). Enlargement of the humeral epicondyle increases the attachment sites of several forelimb muscles (i.e., flexors, pronators, and supinators) needed to generate power and force to dig large burrows (Davis 1964; Hildebrand 1985a; Lessa and Stein 1992; Lagaria and Youlatos 2006). However, complex burrowers displayed the greatest variation in HEI values, suggesting that the relationship between enlarged humeral epicondyles and complex burrowing is not tightly linked. Supporting this idea is the caveat that only four lagomorphs (two pikas and two rabbits) were categorized as complex burrowers. Average HEI values of these two pikas and two rabbits were 0.21 and 0.14, respectively, the latter of which are similar to HEI values from simple and non-burrowers. Correspondingly, the PGLS_ecotype_ model was the equally best supported model for HEI and revealed that the presence of these enlarged humeral epicondyles appear to be driven by pikas (Fig. 7D; Table 5, S5). Under this model, pikas exhibited the highest HEI values, whereas rabbits and hares tended to exhibit similar HEI values. Pikas also tended to display the highest femoral epicondylar (FEI) and olecranon length (OLI) indices compared to rabbits and hares under the best fitting models for both indices (Fig. 7F, 8C; Table 5, S5). Enlargement of the humeral and femoral epicondyles and olecranon process are typically seen as adaptations for fossorial behaviors. However, the majority of pikas do not dig extensive burrows and instead live in crevices found in rocky talus-meadow habitats (Wilson et al. 2016). Therefore, these joint morphologies may not be adaptations to specific burrowing behaviors and instead are simply an ancestral condition in which leporids (hares and rabbits) evolved away from and towards adaptations for more cursorial and saltatorial forms of locomotion. Although (Gambaryan 1974) hypothesized that pikas most closely resembled the ancestral form of lagomorphs and originated as talus-dwellers, the evolutionary history of lagomorphs remains uncertain and requires further investigation to test the evolution and appearance of pika-like and hare/rabbit-like body plans (Ruedas et al. 2018).

## Conclusion

Despite their unique body plans compared to many other terrestrial mammals (Kraatz et al. 2021), lagomorphs exhibit similar patterns in the evolution of their body shapes. Specifically, we found that relative rib length and elongation/shortening of the thoracic and lumbar regions contributed the most to lagomorph body shape evolution, increasingly elongate leporids exhibited relatively shorter limbs, and increasingly larger lagomorphs exhibited relatively more robust body shapes. These body shape patterns largely mirror what has been found in carnivorans and some sciurids (i.e., ground squirrels), the only other mammal clades in which these relationships have been examined (Law et al. 2019; Law 2021b; Linden et al. 2023). Thus, it is tempting to postulate that these body shape patterns may be a universal trend across all mammals. Nevertheless, additional work quantifying body shapes in other mammalian clades is clearly needed to further confirm these patterns.

Lagomorphs, however, differ from the allometric trends in the limbs generally found in other mammals. Whereas many other mammals tend to exhibit relatively more robust limbs with increasing body size (Biewener 1983; Bertram and Biewener 1990; Christiansen 1999; Campione and Evans 2012), extant lagomorphs exhibit increasingly more gracile humeri and ulnae with increasing body size. Cursoriality in larger lagomorphs—and its effects of increasing stride length—is probably what drives divergence from this allometric trend across mammal limbs. Interestingly and contrary to our predictions, we found that models incorporating locomotor and burrowing behaviors were often poor fits to nearly all appendicular functional proxies. These results suggest that lagomorphs display greater variation within each behavioral category than originally predicted and that size dictates appendicular evolution in lagomorphs. Alternatively, our discrete categories may be poor proxies of locomotor and burrowing behaviors. Unfortunately, natural history or performance data remain unavailable for the majority of lagomorphs and prevents us from using finer resolution data such as running speed, leaping abilities, and digging propensity.

Overall, our work reveals unique allometric trends in the appendicular skeleton of lagomorphs whereas the patterns of body shape variation are similar with other studied terrestrial mammals. These patterns demonstrate the complex relationships between morphological evolution, locomotor diversity, and allometry across large macroevolutionary scales. Future work expanding analyses of body plans to other small mammals and “rabbit-like” mammals (e.g., maras and viscachas) will further reveal whether lagomorphs exhibit unique skeletal patterns in their limbs or if there is evidence of skeletal convergence. Altogether, this work provides a strong morphological foundation for future research investigating the evo-devo and macroevolution of mammalian body plans.

## Acknowledgements

The authors are grateful to the collections and its staff of the American Museum of Natural History, Burke Museum of Natural History and Culture at the University of Washington, Field Museum of Natural History, Florida Museum of Natural History, Museum of Vertebrate Zoology at UC Berkeley, University of Alaska Museum of the North, University of Puget Sound Museum, and National Museum of Natural History. We thank Mirra Chinta and Suhyeon Kim for helping collect data, Skipper Lynch for line drawings, and members of the Santana lab at the University of Washington for helpful discussions. The authors were supported by the United States National Science Foundation awards DBI–2128146 and DEB-2447166 (awarded to CJL). N.B. was further supported by a Katharina Casey Leadership Award through the Department of Biology at the University of Washington.

## Notes

### Competing Interest Statement

The authors have declared no competing interest.

