## Supplementary Materials for "Scaling and ecomorphology of lagomorph body shape and appendicular skeleton"

Contains:

**Table S1.** Burrowing behavior categories and references.

**Table S2.** Comparisons of the best-fitting phylogenetic generalized least squares (PGLS) models in body shape and underlying skeletal components.

**Table S3.** The observed difference between the mean coefficients of categorical pairs and its 95% confidence interval.

**Table S4.** Comparisons of the best-fitting phylogenetic generalized least squares (PGLS) models in limb functional proxies.

**Table S5.** The observed difference between the mean coefficients of categorical pairs and its 95% confidence interval.

**References**

**Table S1.** Burrowing behavior categories and references.

| Species | Burrowing category | Justification | Source |
| --- | --- | --- | --- |
| *Brachylagus idahoensis* | Complex burrower | Constructs extensive burrows with several entrances and chambers | Wilson et al. 2016 |
| *Lepus alleni* | Non-burrower | Shelters in forms but no mention of burrowing | Wilson et al. 2016 |
| *Lepus americanus* | Non-burrower | Shelters in forms but no mention of burrowing | Wilson et al. 2016 |
| *Lepus arcticus* | Simple burrower | Digs in snow | Best & Henry 1994 |
| *Lepus californicus* | Simple burrower | Digs burrows deep enough to enclose an individual | Wilson et al. 2016 |
| *Lepus callotis* | Simple burrower | Creates forms up averaging 37 cm in length, 18.3 cm in width, and 6.3 cm in depth | Wilson et al. 2016; Myers et al. 2025 |
| *Lepus capensis* | Simple burrower | Digs short burrows to escape the sun | Wilson et al. 2016 |
| *Lepus europaeus* | Non-burrower | Shelters in forms but no mention of burrowing | Bock 2020 |
| *Lepus flavigularis* | Non-burrower | Shelters in forms but no mention of burrowing | Farias 2004 |
| *Lepus habessinicus* | Non-burrower | Shelters in forms but no mention of burrowing | Wilson et al. 2016; Myers et al. 2025 |
| *Lepus nigricollis* | Non-burrower | Uses burrows or nests created by other species, but no mention of digging | Wilson et al. 2016 |
| *Lepus oiostolus* | Non-burrower | Uses burrows or nests created by other species, but no mention of digging | Myers et al. 2025 |
| *Lepus othus* | Non-burrower | Shelters in forms but no mention of burrowing | Myers et al. 2025 |
| *Lepus peguensis* | Non-burrower | Shelters in forms but no mention of burrowing | Wilson et al. 2016; Myers et al. 2025 |
| *Lepus saxatilis* | Non-burrower | Shelters in forms but no mention of burrowing | Myers et al. 2025 |
| *Lepus tibetanus* | Non-burrower | Shelters in forms but no mention of burrowing | Myers et al. 2025 |
| *Lepus timidus* | Simple burrower | Occasionally digs burrows for young, but does not make extensive use of them | Myers et al. 2025 |
| *Lepus tolai* | Simple burrower | Makes extensive use of shallow depressions that are deeper in windy or cold conditions | Smith, et al. 2010 |
| *Lepus townsendii* | Simple burrower | Makes extensive use of large depressions | Lim 1987 |
| *Nesolagus timminsi* | Non-burrower | Uses burrows or nests created by other species | Myers et al. 2025 |
| *Ochotona alpina* | Non-burrower | Creates nests with grass and hay rather than burrows | Wilson et al. 2016 |
| *Ochotona collaris* | Non-burrower | Uses rock dwellings rather than burrows | Wilson et al. 2016; Myers et al. 2025 |
| *Ochotona curzoniae* | Complex burrower | Creates burrows with several branches and chambers | Wilson et al. 2016 |
| *Ochotona princeps* | Non-burrower | Uses rock dwellings rather than burrows | Markham & Whicker 1972 |
| *Ochotona rufescens* | Complex burrower | Creates burrows with several chambers | Wilson et al. 2016 |
| *Ochotona thibetana* | Simple burrower | Creates burrows | Wilson et al. 2016 |
| *Oryctolagus cuniculus* | Complex burrower | Digs extensive underground burrows for rest and protection | Wilson et al. 2016 |
| *Poelagus marjorita* | Simple burrower | Young are born in short burrows | Wilson et al. 2016; Myers et al. 2025 |
| *Pronolagus crassicaudatus* | Non-burrower | Hides in rock crevices, under boulders, or in dense grass. No mention of burrowing. | [Kovacs & Oroian 2023](https://www.proquest.com/docview/3058285082?sourcetype=Scholarly%20Journals) |
| *Pronolagus rupestris* | Non-burrower | Hides in rock crevices, under boulders, or in dense grass. No mention of burrowing. | Wilson et al. 2016 |
| *Romerolagus diazi* | Simple burrower | Use burrows with hidden entrances | Wilson et al. 2016 |
| *Sylvilagus aquaticus* | Non-burrower | Creates forms but does not make burrows | Wilson et al. 2016; Myers et al. 2025 |
| *Sylvilagus audubonii* | Simple burrower | Creates short burrows for raising young | Chapman & Wilner 1978 |
| *Sylvilagus bachmani* | Non-burrower | Occasionally uses burrows but does not dig its own | Wilson et al. 2016 |
| *Sylvilagus brasiliensis* | Non-burrower | Not known to burrow | Wilson et al. 2016; Myers et al. 2025 |
| *Sylvilagus floridanus* | Non-burrower | Makes nests using grasses | Wilson et al. 2016; Myers et al. 2025 |
| *Sylvilagus nuttallii* | Non-burrower | Occasionally uses burrows but does not dig its own | Wilson et al. 2016 |
| *Sylvilagus palustris* | Simple burrower | Digs holes with long toenails | Chapman & Wilner 1981 |
| *Sylvilagus transitionalis* | Non-burrower | Does not burrow | Wilson et al. 2016 |

**Table S2.** Comparisons of the best-fitting phylogenetic generalized least squares (PGLS) models in body shape and underlying skeletal components. Size-corrected Akaike Information Criterion (AICc) was used to assess model fits. Models where ΔAICc < 2 were considered best supported.

| trait | model | AICc | ΔAICc | AICcW |
| --- | --- | --- | --- | --- |
| hbER |  |  |  |  |
|  | **PGLS_size_** | **-79.74** | **0.00** | **0.96** |
|  | PGLS_ecotype_ | -72.40 | 7.34 | 0.03 |
|  | PGLS_ecotype*size_ | -45.25 | 34.49 | 0.00 |
|  | PGLS_ecotype+size_ | -70.89 | 8.85 | 0.01 |
|  | PGLS_burrow_ | -56.68 | 23.07 | 0.00 |
|  | PGLS_burrow*size_ | -36.36 | 43.38 | 0.00 |
|  | PGLS_burrow+size_ | -59.99 | 19.76 | 0.00 |
|  | PGLS_locomotion_ | -56.72 | 23.02 | 0.00 |
|  | PGLS_locomotion*size_ | -42.47 | 37.27 | 0.00 |
|  | PGLS_locomotion+size_ | -59.88 | 19.86 | 0.00 |
| headER | |  |  |  |
|  | **PGLS_size_** | **-58.31** | **0.00** | **0.74** |
|  | PGLS_ecotype_ | -53.65 | 4.66 | 0.07 |
|  | PGLS_ecotype*size_ | -20.37 | 37.94 | 0.00 |
|  | PGLS_ecotype+size_ | -43.10 | 15.20 | 0.00 |
|  | PGLS_burrow_ | -55.50 | 2.80 | 0.18 |
|  | PGLS_burrow*size_ | -17.72 | 40.59 | 0.00 |
|  | PGLS_burrow+size_ | -44.92 | 13.39 | 0.00 |
|  | PGLS_locomotion_ | -49.43 | 8.88 | 0.01 |
|  | PGLS_locomotion*size_ | -13.70 | 44.61 | 0.00 |
|  | PGLS_locomotion+size_ | -38.43 | 19.88 | 0.00 |
| relative rib length | |  |  |  |
|  | PGLS_size_ | -85.76 | 5.20 | 0.07 |
|  | **PGLS_ecotype_** | **-90.96** | **0.00** | **0.93** |
|  | PGLS_ecotype*size_ | -54.73 | 36.23 | 0.00 |
|  | PGLS_ecotype+size_ | -80.24 | 10.72 | 0.00 |
|  | PGLS_burrow_ | -77.24 | 13.72 | 0.00 |
|  | PGLS_burrow*size_ | -41.55 | 49.41 | 0.00 |
|  | PGLS_burrow+size_ | -66.26 | 24.70 | 0.00 |
|  | PGLS_locomotion_ | -77.19 | 13.77 | 0.00 |
|  | PGLS_locomotion*size_ | -51.44 | 39.52 | 0.00 |
|  | PGLS_locomotion+size_ | -66.22 | 24.74 | 0.00 |
| cervical AEI | |  |  |  |
|  | **PGLS_size_** | **-67.15** | **0.00** | **0.77** |
|  | PGLS_ecotype_ | -64.29 | 2.86 | 0.19 |
|  | PGLS_ecotype*size_ | -26.51 | 40.64 | 0.00 |
|  | PGLS_ecotype+size_ | -53.75 | 13.41 | 0.00 |
|  | PGLS_burrow_ | -54.91 | 12.25 | 0.00 |
|  | PGLS_burrow*size_ | -21.89 | 45.26 | 0.00 |
|  | PGLS_burrow+size_ | -47.39 | 19.76 | 0.00 |
|  | PGLS_locomotion_ | -61.26 | 5.90 | 0.04 |
|  | PGLS_locomotion*size_ | -24.85 | 42.30 | 0.00 |
|  | PGLS_locomotion+size_ | -51.44 | 15.72 | 0.00 |
| thoracic AEI | |  |  |  |
|  | **PGLS_size_** | **-63.84** | **0.00** | **1.00** |
|  | PGLS_ecotype_ | -44.83 | 19.02 | 0.00 |
|  | PGLS_ecotype*size_ | -20.39 | 43.46 | 0.00 |
|  | PGLS_ecotype+size_ | -47.80 | 16.04 | 0.00 |
|  | PGLS_burrow_ | -35.61 | 28.24 | 0.00 |
|  | PGLS_burrow*size_ | -19.04 | 44.81 | 0.00 |
|  | PGLS_burrow+size_ | -46.59 | 17.26 | 0.00 |
|  | PGLS_locomotion_ | -36.96 | 26.88 | 0.00 |
|  | PGLS_locomotion*size_ | -18.68 | 45.17 | 0.00 |
|  | PGLS_locomotion+size_ | -46.40 | 17.45 | 0.00 |
| lumbar AEI | |  |  |  |
|  | **PGLS_size_** | **-35.44** | **1.13** | **0.25** |
|  | **PGLS_ecotype_** | **-36.57** | **0.00** | **0.44** |
|  | PGLS_ecotype*size_ | -8.44 | 28.13 | 0.00 |
|  | **PGLS_ecotype+size_** | **-35.72** | **0.86** | **0.29** |
|  | PGLS_burrow_ | -29.82 | 6.75 | 0.02 |
|  | PGLS_burrow*size_ | -2.53 | 34.04 | 0.00 |
|  | PGLS_burrow+size_ | -24.27 | 12.31 | 0.00 |
|  | PGLS_locomotion_ | -25.18 | 11.39 | 0.00 |
|  | PGLS_locomotion*size_ | 6.07 | 42.64 | 0.00 |
|  | PGLS_locomotion+size_ | -17.02 | 19.56 | 0.00 |
| sacral AEI | |  |  |  |
|  | **PGLS_size_** | **-22.03** | **0.00** | **0.82** |
|  | PGLS_ecotype_ | -18.94 | 3.10 | 0.18 |
|  | PGLS_ecotype*size_ | 18.90 | 40.93 | 0.00 |
|  | PGLS_ecotype+size_ | -8.49 | 13.54 | 0.00 |
|  | PGLS_burrow_ | -3.71 | 18.32 | 0.00 |
|  | PGLS_burrow*size_ | 22.48 | 44.51 | 0.00 |
|  | PGLS_burrow+size_ | -4.01 | 18.02 | 0.00 |
|  | PGLS_locomotion_ | -10.72 | 11.31 | 0.00 |
|  | PGLS_locomotion*size_ | 15.68 | 37.71 | 0.00 |
|  | PGLS_locomotion+size_ | -9.01 | 13.02 | 0.00 |

**Table S3.** The observed difference between the mean coefficients of categorical pairs and its 95% confidence interval. Confidence intervals that do not encompass zero indicates that categorical pairs are significantly different from each other. Bolded cells indicate significant differences between the pair.

| Differences in relative rib length among ecotypes | | |
| --- | --- | --- |
|  | Hare | Pika |
| Rabbit | **0.11 [0.07:0.14]** | **0.14 [0.09:0.19]** |
| Hare |  | 0.03 [-0.08:0.02] |
| Differences in lumbar AEI among ecotypes | | |
|  | Hare | Pika |
| Rabbit | **-0.10 [-0.11:-0.10]** | **-0.40 [-0.40:-0.39]** |
| Hare |  | **0.29 [0.29:0.30]** |

**Table S4.** Comparisons of the best-fitting phylogenetic generalized least squares (PGLS) models in limb functional proxies. Size-corrected Akaike Information Criterion (AICc) was used to assess model fits. Models where ΔAICc < 2 were considered best supported.

| Limb indices | model | AICc | ΔAICc | AICcW |
| --- | --- | --- | --- | --- |
| Scapula index (SI) | |  |  |  |
|  | **PGLS_size_** | **-6.67** | **0.00** | **0.84** |
|  | PGLS_ecotype_ | -0.18 | 6.50 | 0.03 |
|  | PGLS_ecotype*size_ | 32.44 | 39.12 | 0.00 |
|  | PGLS_ecotype+size_ | 6.71 | 13.38 | 0.00 |
|  | PGLS_burrow_ | 0.23 | 6.90 | 0.03 |
|  | PGLS_burrow*size_ | 35.27 | 41.95 | 0.00 |
|  | PGLS_burrow+size_ | 10.93 | 17.61 | 0.00 |
|  | PGLS_locomotion_ | -2.38 | 4.30 | 0.10 |
|  | PGLS_locomotion*size_ | 34.70 | 41.38 | 0.00 |
|  | PGLS_locomotion+size_ | 6.86 | 13.53 | 0.00 |
| Brachial index (BI) | |  |  |  |
|  | **PGLS_size_** | **-71.65** | **0.62** | **0.41** |
|  | **PGLS_ecotype_** | **-72.26** | **0.00** | **0.55** |
|  | PGLS_ecotype*size_ | -33.91 | 38.36 | 0.00 |
|  | PGLS_ecotype+size_ | -61.64 | 10.62 | 0.00 |
|  | PGLS_burrow_ | -62.07 | 10.19 | 0.00 |
|  | PGLS_burrow*size_ | -26.90 | 45.37 | 0.00 |
|  | PGLS_burrow+size_ | -52.86 | 19.41 | 0.00 |
|  | PGLS_locomotion_ | -66.34 | 5.92 | 0.03 |
|  | PGLS_locomotion*size_ | -35.90 | 36.37 | 0.00 |
|  | PGLS_locomotion+size_ | -63.53 | 8.73 | 0.01 |
| Humeral Robustness Index (HRI) | |  |  |  |
|  | **PGLS_size_** | **-262.85** | **0.00** | **0.87** |
|  | PGLS_ecotype_ | -258.46 | 4.39 | 0.10 |
|  | PGLS_ecotype*size_ | -225.45 | 37.40 | 0.00 |
|  | PGLS_ecotype+size_ | -247.80 | 15.05 | 0.00 |
|  | PGLS_burrow_ | -253.29 | 9.56 | 0.01 |
|  | PGLS_burrow*size_ | -218.70 | 44.15 | 0.00 |
|  | PGLS_burrow+size_ | -246.23 | 16.62 | 0.00 |
|  | PGLS_locomotion_ | -255.44 | 7.41 | 0.02 |
|  | PGLS_locomotion*size_ | -218.09 | 44.76 | 0.00 |
|  | PGLS_locomotion+size_ | -245.21 | 17.64 | 0.00 |
| Humeral Epicondylar Index (HEI) | |  |  |  |
|  | PGLS_size_ | -206.46 | 2.95 | 0.13 |
|  | **PGLS_ecotype_** | **-208.35** | **1.06** | **0.32** |
|  | PGLS_ecotype*size_ | -178.16 | 31.24 | 0.00 |
|  | PGLS_ecotype+size_ | -197.74 | 11.66 | 0.00 |
|  | **PGLS_burrow_** | **-209.40** | **0.00** | **0.55** |
|  | PGLS_burrow*size_ | -176.24 | 33.17 | 0.00 |
|  | PGLS_burrow+size_ | -198.42 | 10.98 | 0.00 |
|  | PGLS_locomotion_ | -198.82 | 10.59 | 0.00 |
|  | PGLS_locomotion*size_ | -160.22 | 49.18 | 0.00 |
|  | PGLS_locomotion+size_ | -187.82 | 21.58 | 0.00 |
| Olecranon Length Index (OLI) | |  |  |  |
|  | **PGLS_size_** | **-213.30** | **1.94** | **0.27** |
|  | **PGLS_ecotype_** | **-215.24** | **0.00** | **0.71** |
|  | PGLS_ecotype*size_ | -180.66 | 34.59 | 0.00 |
|  | PGLS_ecotype+size_ | -205.29 | 9.96 | 0.01 |
|  | PGLS_burrow_ | -206.11 | 9.14 | 0.01 |
|  | PGLS_burrow*size_ | -168.42 | 46.83 | 0.00 |
|  | PGLS_burrow+size_ | -195.11 | 20.13 | 0.00 |
|  | PGLS_locomotion_ | -207.21 | 8.03 | 0.01 |
|  | PGLS_locomotion*size_ | -171.48 | 43.76 | 0.00 |
|  | PGLS_locomotion+size_ | -198.52 | 16.73 | 0.00 |
| Ulna Robustness Index (URI) | |  |  |  |
|  | **PGLS_size_** | **-231.13** | **0.00** | **0.93** |
|  | PGLS_ecotype_ | -224.68 | 6.45 | 0.04 |
|  | PGLS_ecotype*size_ | -186.63 | 44.50 | 0.00 |
|  | PGLS_ecotype+size_ | -214.26 | 16.87 | 0.00 |
|  | PGLS_burrow_ | -219.18 | 11.95 | 0.00 |
|  | PGLS_burrow*size_ | -186.72 | 44.41 | 0.00 |
|  | PGLS_burrow+size_ | -212.27 | 18.86 | 0.00 |
|  | PGLS_locomotion_ | -224.19 | 6.94 | 0.03 |
|  | PGLS_locomotion*size_ | -190.55 | 40.58 | 0.00 |
|  | PGLS_locomotion+size_ | -213.29 | 17.84 | 0.00 |
| Manus Proportion Index (MANUS) | |  |  |  |
|  | **PGLS_size_** | **-168.91** | **0.00** | **0.78** |
|  | PGLS_ecotype_ | -166.15 | 2.76 | 0.20 |
|  | PGLS_ecotype*size_ | -135.00 | 33.90 | 0.00 |
|  | PGLS_ecotype+size_ | -158.87 | 10.04 | 0.01 |
|  | PGLS_burrow_ | -160.55 | 8.36 | 0.01 |
|  | PGLS_burrow*size_ | -125.86 | 43.04 | 0.00 |
|  | PGLS_burrow+size_ | -149.77 | 19.13 | 0.00 |
|  | PGLS_locomotion_ | -160.41 | 8.50 | 0.01 |
|  | PGLS_locomotion*size_ | -131.12 | 37.79 | 0.00 |
|  | PGLS_locomotion+size_ | -149.93 | 18.98 | 0.00 |
| Crural Index (Cl) | |  |  |  |
|  | PGLS_size_ | -90.50 | 3.23 | 0.14 |
|  | **PGLS_ecotype_** | **-93.74** | **0.00** | **0.71** |
|  | PGLS_ecotype*size_ | -60.37 | 33.36 | 0.00 |
|  | PGLS_ecotype+size_ | -88.14 | 5.60 | 0.04 |
|  | PGLS_burrow_ | -82.15 | 11.59 | 0.00 |
|  | PGLS_burrow*size_ | -49.79 | 43.95 | 0.00 |
|  | PGLS_burrow+size_ | -71.63 | 22.11 | 0.00 |
|  | PGLS_locomotion_ | -89.99 | 3.75 | 0.11 |
|  | PGLS_locomotion*size_ | -54.01 | 39.73 | 0.00 |
|  | PGLS_locomotion+size_ | -81.04 | 12.70 | 0.00 |
| Femoral Robustness Index (FRI) | |  |  |  |
|  | **PGLS_size_** | **-261.77** | **0.00** | **0.81** |
|  | PGLS_ecotype_ | -255.50 | 6.27 | 0.04 |
|  | PGLS_ecotype*size_ | -220.50 | 41.27 | 0.00 |
|  | PGLS_ecotype+size_ | -245.33 | 16.44 | 0.00 |
|  | PGLS_burrow_ | -258.33 | 3.43 | 0.15 |
|  | PGLS_burrow*size_ | -222.29 | 39.48 | 0.00 |
|  | PGLS_burrow+size_ | -247.94 | 13.83 | 0.00 |
|  | PGLS_locomotion_ | -252.91 | 8.86 | 0.01 |
|  | PGLS_locomotion*size_ | -215.04 | 46.73 | 0.00 |
|  | PGLS_locomotion+size_ | -242.32 | 19.45 | 0.00 |
| Femoral Epicondylar Index (FEl) | |  |  |  |
|  | **PGLS_size_** | **-213.52** | **0.00** | **0.69** |
|  | **PGLS_ecotype_** | **-211.82** | **1.70** | **0.30** |
|  | PGLS_ecotype*size_ | -177.42 | 36.10 | 0.00 |
|  | PGLS_ecotype+size_ | -203.54 | 9.98 | 0.01 |
|  | PGLS_burrow_ | -202.47 | 11.05 | 0.00 |
|  | PGLS_burrow*size_ | -170.38 | 43.14 | 0.00 |
|  | PGLS_burrow+size_ | -193.85 | 19.67 | 0.00 |
|  | PGLS_locomotion_ | -204.38 | 9.14 | 0.01 |
|  | PGLS_locomotion*size_ | -169.01 | 44.51 | 0.00 |
|  | PGLS_locomotion+size_ | -195.05 | 18.47 | 0.00 |
| Tibial Robustness Index (TRI) | |  |  |  |
|  | **PGLS_size_** | **-234.40** | **0.00** | **0.89** |
|  | PGLS_ecotype_ | -229.65 | 4.75 | 0.08 |
|  | PGLS_ecotype*size_ | -205.75 | 28.64 | 0.00 |
|  | PGLS_ecotype+size_ | -224.61 | 9.79 | 0.01 |
|  | PGLS_burrow_ | -224.88 | 9.52 | 0.01 |
|  | PGLS_burrow*size_ | -190.35 | 44.04 | 0.00 |
|  | PGLS_burrow+size_ | -216.11 | 18.28 | 0.00 |
|  | PGLS_locomotion_ | -226.50 | 7.90 | 0.02 |
|  | PGLS_locomotion*size_ | -187.96 | 46.44 | 0.00 |
|  | PGLS_locomotion+size_ | -214.63 | 19.76 | 0.00 |
| Pes Proportion Index (PES) | |  |  |  |
|  | **PGLS_size_** | **-130.27** | **1.06** | **0.36** |
|  | **PGLS_ecotype_** | **-131.32** | **0.00** | **0.62** |
|  | PGLS_ecotype*size_ | -95.07 | 36.26 | 0.00 |
|  | PGLS_ecotype+size_ | -123.04 | 8.29 | 0.01 |
|  | PGLS_burrow_ | -121.31 | 10.02 | 0.00 |
|  | PGLS_burrow*size_ | -86.09 | 45.24 | 0.00 |
|  | PGLS_burrow+size_ | -110.59 | 20.73 | 0.00 |
|  | PGLS_locomotion_ | -121.63 | 9.69 | 0.01 |
|  | PGLS_locomotion*size_ | -82.98 | 48.34 | 0.00 |
|  | PGLS_locomotion+size_ | -110.88 | 20.45 | 0.00 |
| Intermembral Index (IM) | |  |  |  |
|  | **PGLS_size_** | **-118.31** | **0.00** | **0.81** |
|  | PGLS_ecotype_ | -115.25 | 3.07 | 0.18 |
|  | PGLS_ecotype*size_ | -92.29 | 26.02 | 0.00 |
|  | PGLS_ecotype+size_ | -109.24 | 9.07 | 0.01 |
|  | PGLS_burrow_ | -103.49 | 14.83 | 0.00 |
|  | PGLS_burrow*size_ | -70.71 | 47.61 | 0.00 |
|  | PGLS_burrow+size_ | -98.69 | 19.62 | 0.00 |
|  | PGLS_locomotion_ | -104.13 | 14.18 | 0.00 |
|  | PGLS_locomotion*size_ | -84.07 | 34.24 | 0.00 |
|  | PGLS_locomotion+size_ | -99.78 | 18.54 | 0.00 |

**Table S5.** The observed difference between the mean coefficients of categorical pairs and its 95% confidence interval. Confidence intervals that do not encompass zero indicates that categorical pairs are significantly different from each other. Bolded cells indicate significant differences between the pair.

| Differences in BI among ecotypes | |  |
| --- | --- | --- |
|  | Hare | Pika |
| Rabbit | **0.18 [0.14:0.22]** | **0.13 [0.20: 0.07]** |
| Hare |  | **0.32 [0.25:0.38]** |
| Differences in HEI among burrowing behaviors | | |
|  | Complex | None |
| Simple | **0.03 [0.03:0.03]** | -0.00 [-0.00: 0.00] |
| Complex |  | **0.03 [0.03:0.03]** |
| Differences in HEI among ecotypes | | |
|  | Hare | Pika |
| Rabbit | **-0.01 [-0.01:-0.01]** | **0.05 [0.05:0.05]** |
| Hare |  | **-0.06 [-0.06:-0.06]** |
| Differences in OLI among ecotypes | | |
|  | Hare | Pika |
| Rabbit | **0.02 [0.03:0.01]** | **0.02 [0.01:0.03]** |
| Hare |  | **0.04 [0.03:0.05]** |
| Differences in CI among ecotypes | |  |
|  | Hare | Pika |
| Rabbit | **0.08 [0.04:0.12]** | 0.04 [-0.01:0.09] |
| Hare |  | 0.05 [-0.01:0.09] |
| Differences in PES among ecotypes | | |
|  | Hare | Pika |
| Rabbit | **-0.00 [-0.00:-0.00]** | **0.02 [0.02:0.02]** |
| Hare |  | **-0.03 [-0.03:-0.03]** |
| Differences in PES among ecotypes | | |
|  | Hare | Pika |
| Rabbit | **0.04 [0.02:0.06]** | **0.04 [0.01:0.07]** |
| Hare |  | **0.08 [0.05:0.11]** |
